# Stress Induced Signaling Pathways in Burkitt’s Lymphoma Play Novel Mechanisms in Fate Determination and Pathogenesis of Germinal Center-Derived B-Lymphomas

**DOI:** 10.1101/2024.12.19.628635

**Authors:** Santosh K Gothwal, Jacqueline H Barlow

## Abstract

B cell receptor signaling, NF-κB signaling, BCL6 and p53 regulation are essential for germinal center (GC) B cell fate. Dysregulation of these pathways drives the pathogenesis and treatment resistance of GC-derived B-lymphomas (GCDBL). To explore how these pathways affect GCDBL fate and pathogenesis, we treated Raji cells (a GCDBL and Burkitt’s lymphoma) with mild hydroxyurea (HU) to simulate genotoxic stress encountered by GC B cells. Genome-wide mapping of histone H3K4me3 and p53 target analysis in HU-treated Raji cells combined with transcriptome analysis of human tonsil GC B cells identified *ATAD2B* (a p53 target) as differentially expressed. We found that p53 suppresses *ATAD2B* and *ATAD2*, while *ATAD2* and *BCL6* transcripts positively correlate in DLBCLs, suggesting that p53 regulates BCL6 in GC B cells via ATAD2 suppression. We propose that p53 regulation of *BCL6* quality assures GC B cells before GC exit. Unlike BCL6 suppression of IFN-γ and NF-κB signaling in GC B cells, we identified *IFNGR1* as a loosely bound BCL6 target and observed loss of BCL6 regulation on genes encoding inhibitory subunits of NF-κB signaling in B-lymphoma treated with a Bruton tyrosine kinase (BTK) inhibitor. These adaptations, alongside with prevalent genetic inactivation of NF-κB inhibitory genes in DLBLCs, likely contribute to DLBCL pathogenesis and therapy resistance. Our findings highlight the pivotal role of the p53-BCL6 axis in GC B cell fate and its dysregulation in DLBCL pathogenesis and chemoresistance.

**Key Highlights:**

1. Genes induced in human Burkitt’s lymphoma under genotoxic stress are largely independent of histone H3K4me3 marks at transcriptional start sites (TSS).
2. Loss of BCL6 regulation on genes encoding components of IFN-γ signaling is associated with reduced survival and the pathogenesis of DLBCL.
3. Inactivating mutations in genes encoding components of the NF-κB inhibitory subunit serve as an adaptation for DLBCL pathogenesis.
4. *BCL6* expression is correlated with *ATAD2* overexpression in DLBCL and solid tumors.

## Introduction

The germinal center (GC) reaction is crucial for the development of high-affinity B cells, primarily mediated by activation-induced cytidine deaminase (AID). AID initiates DNA double-strand breaks (DSBs) at the immunoglobulin (*IG*) locus of GC B cells, leading to somatic hypermutation (SHM) and class switch recombination (CSR), essential processes for producing high-affinity antibodies [1–4]. However, AID off-target activity can result in chromosomal translocations, creating oncogenic precursors of B-lymphomas referred to as GCDBLs [5]. Notable translocations, such as *IGH-MYC*, characteristic of Burkitt’s lymphoma, and *IG-BCL6*, observed in diffuse large B cell lymphoma (DLBCL), pose significant risks during the GC reaction [6–10]. Within the differentiation pathways of GC B cells, GCDBLs can evolve into memory B cells or plasma cells, leading to conditions like B cell chronic lymphocytic leukemia (B-CLL) or multiple myeloma (MM), respectively [11]. Conversely, B-lymphomas emerging from dark zone (DZ) and light zones (LZ) can be matured into DLBCLs and follicular lymphomas. For effective GC outcomes, it is essential to promote the survival of GC B cells that undergo error-free SHM and CSR, generating high-affinity B cell receptors (BCRs) [12]. Simultaneously, it is critical to suppress the survival and differentiation of GCDBL precursors, such as those harboring *IGH-MYC* translocations, which are a hallmark of Burkitt’s lymphoma [13]. The pathways regulating these processes remain unknown.

BCL6 plays a pivotal role in protecting GC B cells from genotoxic stress-induced apoptotic signals present in the GC microenvironment [14]. As a transcriptional repressor, BCL6 inhibits genes involved in the DNA damage response, p53 regulation, inflammatory signaling, and various cell death pathways [15, 16]. This repression allows GC B cells to evade apoptosis until they acquire high-affinity BCRs and differentiate into memory or plasma cells[15, 16]. The dynamics of BCR signaling and NF-κB signaling are integral to the activation, survival, and differentiation of GC B cells. Notably, BCL6 suppresses both BCR and NF-κB signaling to ensure GC B cell viability. Additionally, G-protein-coupled receptors (GPCRs), such as CXCR4, CXCR5, CCR7, and CD86, along with associated G-proteins, facilitate the signaling, migration, and survival of GC B cells as they navigate the distinct zones of the GC such as the DZ, LZ, and the T-B border [17–19]. The survival of GC B cells in the LZ is also supported by their capacity for antigen presentation, maintained by CREBBP and CIITA, ensuring T cell-mediated help and feedback from follicular dendritic cells (FDCs) [20–23].

Upregulation of BCR, NF-κB, and GPCR signaling, along with alterations in p53 and G-protein signaling, have been linked to B-lymphoma pathogenesis and therapeutic resistance [24, 25]. However, how modifications in these pathways along with the loss of BCL6 regulation and changes in p53 function affect the fate of GCDBLs within the GC microenvironment remains poorly understood. Additionally, whether genetic alterations in genes encoding the core components of NF-κB and IFN-γ signaling confer adaptation and survival advantages that drive B-lymphoma pathogenesis is not well characterized. Addressing these gaps could yield critical insights into the mechanisms underlying both B-lymphoma pathogenesis and treatment resistance. To address these questions, we utilized Raji cells, a Burkitt’s lymphoma cell line with the *IGH-MYC* translocation, as a model for GCDBLs. We induced mild genotoxic stress in Raji cells using hydroxyurea (HU) to simulate the stress experienced by GC B cells, followed by an analysis of differentially expressed genes. By mapping the transcriptome and conducting genome-wide histone H3K4-me3 analysis in HU-treated Raji cells, we identified alterations that influence NF-κB signaling in GCDBLs. In contrast, analysis of datasets of DLBCLs treated with FX1, a BCL6 inhibitor, and ibrutinib, a Bruton tyrosine kinase (BTK) inhibitor, showed decreased expression of genes encoding NF-κB inhibitory subunits, indicating a hijacked regulation of NF-κB signaling that contributes to therapy resistance. Furthermore, p53 target gene analysis in HU-treated Raji cells revealed a novel role for p53 in suppressing ATAD2-dependent *BCL6* expression, correlating with *BCL6* and *ATAD2* expression in DLBCL. These insights may have therapeutic implications not only for B-lymphomas but also for solid tumors.

## Results

### Genome-wide Mapping of Histone H3K4me3 and Transcriptome analysis in HU-treated Raji Cells

To investigate the alteration in molecular pathways influencing GCDBL fate, we used Raji cells as a model cell line (Figure 1A). Similar to GC B cells, Raji cells exhibit CXCR4 and CXCR5 expression (Supplementary Figure 1A) [26, 27], while harboring the AID induced Burkitt’s lymphoma translocation t(8:14) between the immunoglobulin and *MYC* loci (Supplementary Figure 1B) [6, 7, 22, 28]. These features make Raji cells a good model to investigate the mechanisms determining the fate of GCDBLs within the GC microenvironment (Supplementary Figure 1A, B). We hypothesized that mapping transcriptional changes caused by HU induced genotoxic stress in G1-S phase, similar to that experienced by GC B cells during CSR and SHM, could reveal potential alterations in gene expression of components regulating BCR signaling, NF-κB signaling, and BCL6-dependent genes, impacting GC B cell fate [10, 29–33]. We treated Raji cells with 4 mM HU to induce genotoxic stress and measured the differentially expressed genes (DEGs) by RNA-sequencing (Figure 1A-B). To ensure the genes identified were specific to HU-induced stress, we further included groups of Raji cells treated with thymidine, followed by nocodazole (Thy-Noc), which enables synchronization in G2-M phase (Supplementary Figure 1C).

**Figure 1.**
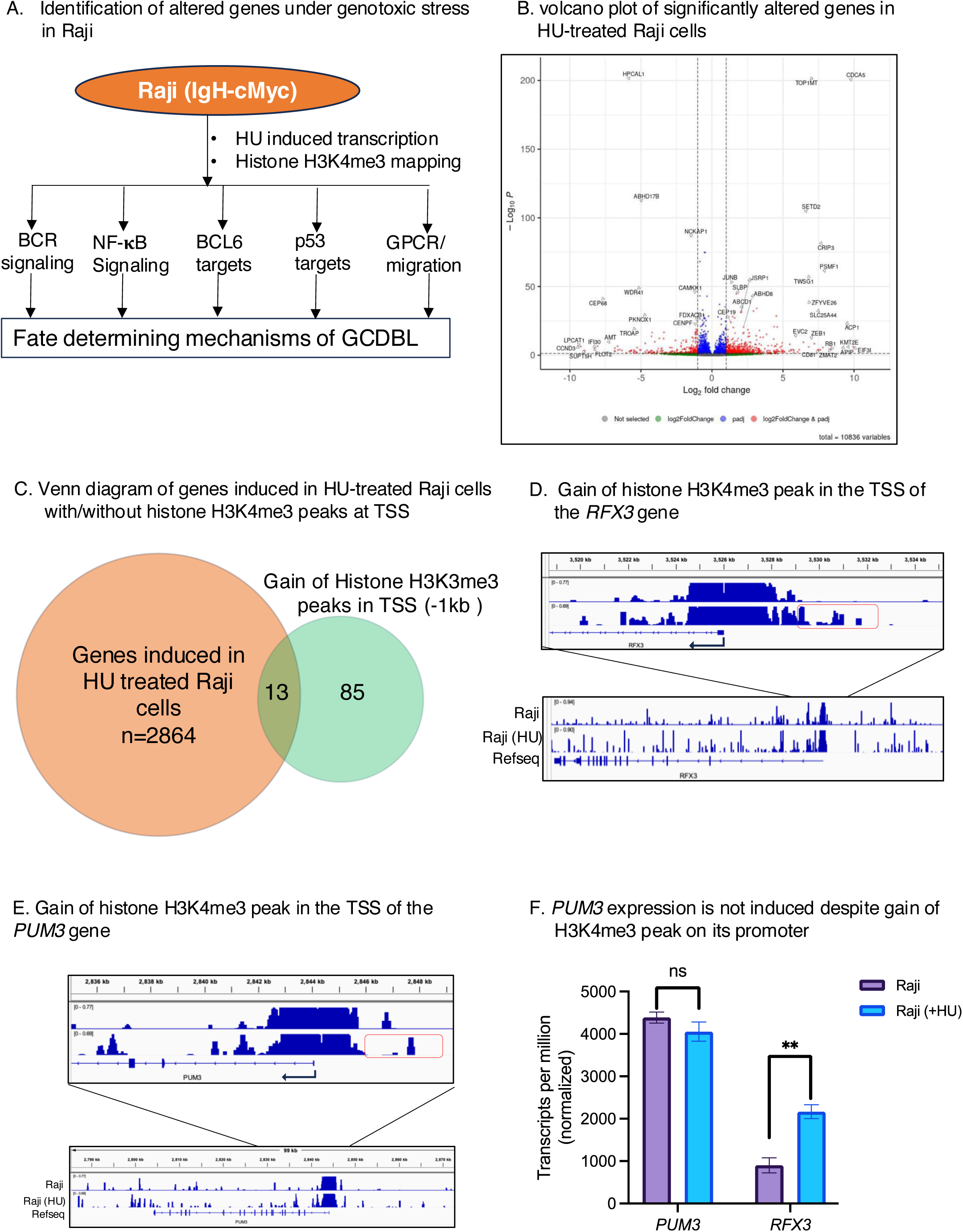
Differentially expressed Genes in HU-treated Raji human Burkitt’s lymphoma cells. **(A)** Schematic representation of HU treatment in Raji cells, inducing transient genotoxic stress, followed by histone H3K4me3 mapping and the identification of differentially expressed genes involved in BCL6 regulation, p53 regulation, BCR signaling, NF-κB signaling, GPCR, and G-protein pathways. **(B)** Volcano plot showing differentially expressed genes between control and HU-treated Raji cells. Differences in gene expression with a cutoff p-value of less than 0.05 are considered statistically significant. The X-axis represents log2 fold changes, and the y-axis represents adjusted p values. Data represent the cumulative results from three independent groups. **(C)** Venn diagram illustrating the relationship between genes induced in RNA-seq and those gaining histone H3K4me3 peaks within the −1 kb region of transcription start sites (TSS) in HU-treated Raji cells. The circles are not drawn to scale. A total of 2864 genes were identified as induced in the RNA-seq analysis with significant p-values < 0.05. Of these, 85 genes showed H3K4me3 peaks within −1 kb of their TSS, and 13 of these 85 genes showed increased expression (log2FC >1). The orange circle represents genes induced in HU-treated Raji cells, while the light green circle represents number of genes whose TSS gained H3K4me3 peaks. **(D)** The increase in histone H3K4me3 peaks at TSSs is not directly correlated with gene expression induced in HU-treated Burkitt’s lymphoma cells. Visualization of histone H3K4me3 peaks in control and HU-treated Raji cells using IGV along the *RFX3* gene. The lower panel displays histone H3K4me3 signal across the *RFX3* gene body and the upper panel shows ChIP-seq signals of histone H3K4me3 peaks near the TSS in HU-treated Raji cells. The red rectangle highlights the gained H3K4me3 signal near the TSS. The scale above the ChIP-seq track represents a 1 kb distance between two consecutive vertical lines. **(E)** Visualization of histone H3K4me3 peaks in control and HU-treated Raji cells, focusing on the *PUM3* gene. The lower panel shows histone H3K4me3 signal across the *PUM3* gene body and the upper panel illustrates ChIP-seq signals of H3K4me3 peaks near the TSS in HU-treated cells. The red rectangle highlights the gained H3K4me3 signal near the TSS. The scale above the ChIP-seq track represents a 1 kb distance between two consecutive vertical lines. **(F)** Normalized counts per million (CPM) values of *PUM3* and *RFX3* in control and HU-treated Raji cells. *PUM3* expression remained unchanged, while *RFX3* expression was significantly induced in HU-treated Raji cells. Unpaired two-tailed t-test: *PUM3*, p > 0.9999 for Raji vs. Raji (HU); *RFX3*, p = 0.0064 for Raji vs. Raji (HU). Data are shown as mean ± SEM, n = 3.

Among the top 50 upregulated genes were *ACP1*, which encodes a small phosphatase (Figure 1B, Supplementary Figure 1D). Comparing the RNA-seq data from normal activated GC B cells (AGCBs) derived from tonsils[34], we observed that *ACP1* expression was high in the light zone (LZ) and plasma cell clusters of AGCBs (Supplementary Figure 1D). Importantly, the role of ACP1 in GC B cells is unknown. Given a differential expression of *ACP1* in HU-treated Raji cells and its association with specific GC compartments in human GCs (Figure 1B, Supplementary Figure 1D), these results suggest a possible role of ACP1 in GC B cells. Zinc finger proteins were another prominent class of genes which were induced by HU treatment (Figure 1B, Supplementary Figure 1D). Notably, *ZNF438*, *ZEB1*, and *ZNF563* were among the top 50 upregulated genes. Interestingly, *ZNF438* and *ZNF563* were not expressed in any subpopulation of AGCBs[34], suggesting that their altered expression could be critical for the fate of GC B cells. *SETD2*, a gene encoding for a histone H3K36-methyltransferase, which plays a critical role in the transcriptional regulation, was also induced in HU-treated Raji cells (Figure 1B, Supplementary Figure 1E). Given the role of SETD2 in transcriptional regulation [35], we confirmed whether Raji cells encode a functional SETD2 proteins. The knockdown by short hairpin RNA (shRNA) targeting *SETD2* in Raji cells reduced the histone H3K36me3 signals compared to scramble knockdown Raji cells (shSCR-Raji) (Supplementary Figure 1F). These results suggest that the DEGs observed in HU-treated Raji cells result from *SETD2* dependent Histone H3K36me3 (Supplementary Figure 1F). On the other hand, top 50 downregulated genes included *ABHD17B*, a positive regulator of N-Ras signaling, a growth stimulator, associated with acute myeloid leukemia pathogenesis (Supplementary Figure 1E)[36]. The downregulation of *ABHD17B* might impair N-Ras signaling in the GCDBL, thereby reducing their survival signals in the GC microenvironment, leading to GCDBL elimination. Notably, *ABHD17B* expression was absent in AGCBs[34], indicating its expression is specific to GCDBLs and may affect the selection of GCDBLs in the GC microenvironment. These results suggest that few genes induced in the HU-treated Raji cells are unique and not expressed in AGCBs, suggesting their expression levels could potentially affect the GCDBL fate.

To know how many genes induced upon HU-treatment in Raji cells are associated with open chromatin at their promoters, we performed genome-wide mapping of histone H3K4me3 mark in control and HU-treated cells (Figure 1C-F). Histone H3K4me3 enrichment at gene promoters strongly correlates with transcriptional activation [37–39]. Our analysis revealed that 85 genes gained H3K4me3 marks within 1 kb upstream of their transcription start sites (TSS), while only 13 genes exhibited induced expression (Log2FC >1) (Figure 1C). However, the expression of most of these genes, including *PUM3,* which is involved in mRNA processing [40], was not significantly altered (Figure 1C-F). This suggests that the increased levels of histone H3K4me3 does not fundamentally induce the gene expression in GCDBLs under genotoxic stress (Figure 1C-F). These results highlight that the dynamics of altered transcription in GC B cells is not strictly coupled to changes in histone H3K4me3, which is consistent with functioning of master regulators such as BCL6 in GC B cells [14]. However, it is noteworthy that most genes exhibiting unaltered H3K4me3 levels in the HU-treated Raji cells already harbor histone H3K4me3 peak in gene promoters indicating these gene promoters have constitutively open chromatin state (Figure 1C-F). Conversely, genes like *RF3* showed increased expression in HU-treated Raji cells (Figure 1D, F), exhibited a gain of H3K4me3 peak in its TSS (Figure 1D), indicating that H3K4me3 enrichment may correlate with expression of a few genes. These results imply a non-essential role for an increase in histone H3K4me3 to induce gene expression in GCDBLs and their induction could be determined by removal of other master transcriptional regulators such BCL6.

### Induced *NFKBIE* expression in HU-treated Raji cells and implication with B-lymphoma pathogenesis

Among the DEGs of HU-treated Raji cells, we next plotted the genes encoding for components regulating BCR and NF-κB signaling in HU-treated Raji cells (Figure 2A, B). BCR signaling plays an essential role in GC B cell survival and differentiation, with key components such as *CD79A*, *CD79B*, *CARD11*, and *LYN* playing essential roles in this process [29–32]. In HU-treated Raji cells, as well as Thy-Noc-treated Raji cells, the expression of BCR signaling components remained unaltered (Figure 2A). This suggests that expression of BCR signaling is not altered at transcriptional level in GCDBL and may not directly influence the fate of GCDBLs emerged having oncogenic translocations. Nevertheless, this does not exclude the possibility that compromised BCR signaling could still affect GCDBL survival by downstream signaling targets, independently of the transcriptional regulation of its core components.

**Figure 2:**
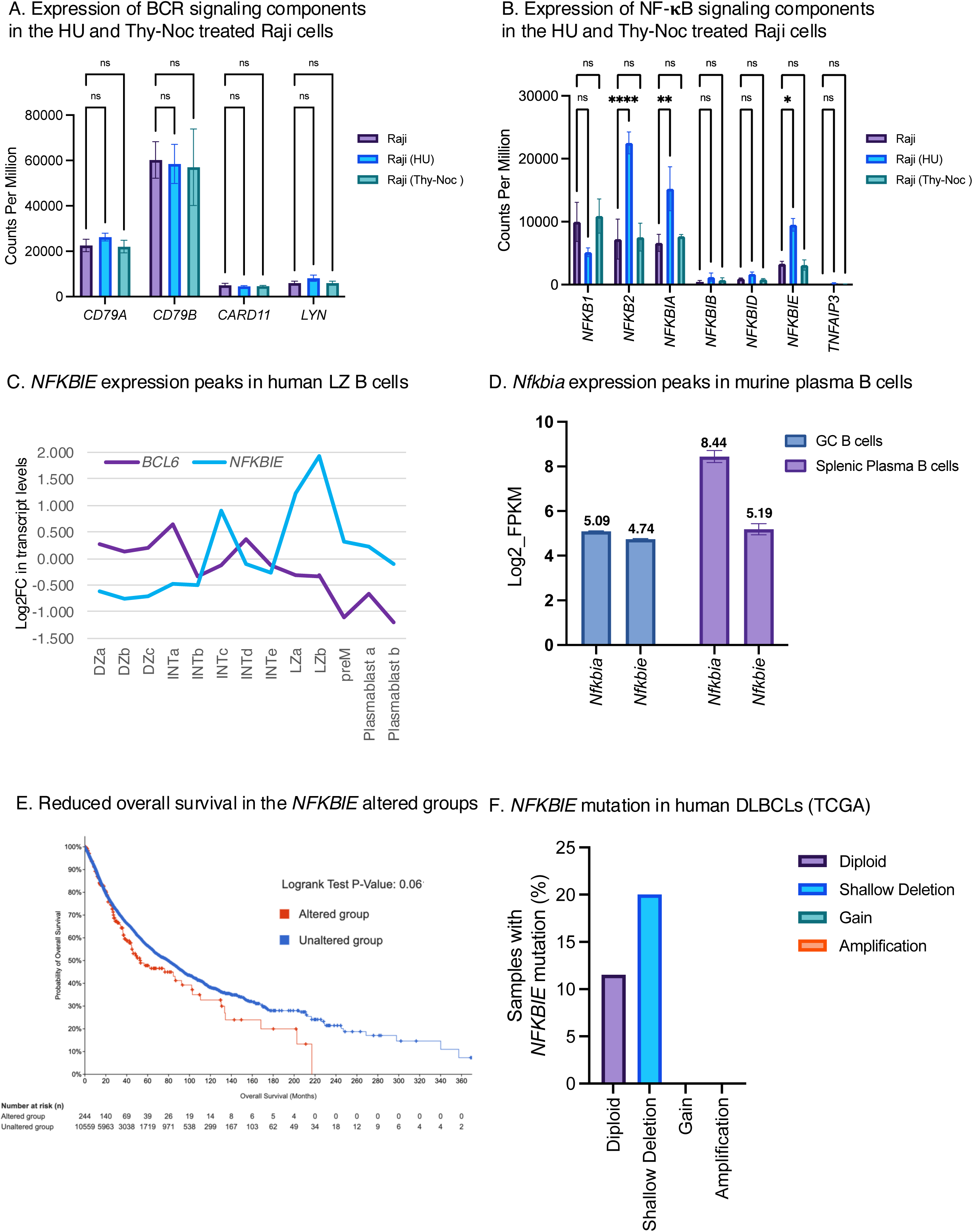
Induced expression of genes encoding the inhibitory subunits of NF-κB signaling in Raji cells. **(A)** Transcript levels of *CD79A*, *CD79B*, *CARD11*, and *LYN* were not altered in Raji (HU) and Raji (Thy-Noc) cells compared to Raji cells. Data are mean ± SEM (gene count values) of three independent samples. **(B)** Transcript levels of *NFKB1*, *NFKB2*, *NFKBIA*, *NFKBIB*, *NFKBID*, *NFKBIE*, and *TNFAIP3* in Raji, Raji (HU), and Raji (Thy-Noc)-treated cells. *NFKB1*; p=0.0892 for Raji vs Raji (HU) and p=0.9081 for Raji vs Raji (Thy-Noc). *NFKB2*; p<0.0001 for Raji vs Raji (HU) and p=0.09895 for Raji vs Raji (Thy-Noc). *NFKBIA*; p=0.0011 for Raji vs Raji (HU) and p=0.8835 for Raji vs Raji (Thy-Noc). *NFKBIB*; p=0.9370 for Raji vs Raji (HU) and p=0.9906 for Raji vs Raji (Thy-Noc). *NFKBID*; p=0.9416 for Raji vs Raji (HU) and p=0.9977 for Raji vs Raji (Thy-Noc). *NFKBIE*; p=0.0216 for Raji vs Raji (HU) and p=0.9959 for Raji vs Raji (Thy-Noc). *TNFAIP3*; p=0.9953 for Raji vs Raji (HU) and p=0.9992 for Raji vs Raji (Thy-Noc). Tukey’s multiple comparison test. Data are presented as mean ± SEM, n=3. **(C)** Correlation analysis of *NFKBIE* and *BCL6* in the DZ sub-populations (DZa, DZb, DZc), intermediate zone subpopulations (INTa, INTb, INTc, INTd, INTe), LZ sub-populations (LZa, LZb), pre-memory (PreM), and plasmablast (plasmablast a, plasmablast b) subpopulations [34]. Log2 fold change values of each transcript are shown in different subpopulations. **(D)** *Nfkbia* and *Nfkbie* expression in mouse splenic B cells, highlighting the increased expression of *Nfkbia* in splenic plasma B cells compared to GC B cells [41] (n = 2 for GC B cells, n = 3 for splenic plasma B cell samples; GSE60927). **(E)** TCGA datasets showing the overall survival of *NFKBIE*-altered groups compared to unaltered groups. Comparison of survival kinetics initiated with 244 altered groups and 10,559 unaltered groups on day 0. The log-rank test was used to assess the statistical difference between the survival distribution (p=0.06) **(F)** Classification of *NFKBIE* mutations in DLBCL samples from TCGA database. Among the 37 profiled DLBCL samples, 10.8% exhibited *NFKBIE* mutations. Of the 26 diploid DLBCL cases, 11.5% showed *NFKBIE* mutations. The observed mutations included frameshift (fs) (*NFKBIE*:Y254Sfs, encoding for 13 amino acids; *NFKBIE*:*L168Pfs* encoding for 42 amino acids), point mutations (*NFKBIE*:*L313Q*, *NFKBIE*:*L441P*, *NFKBIE*:*A251T*), and splicing mutation (X261_splice, A251T).

We next examined the expression of NF-κB signaling components (Figure 2B). NF-κB signaling plays an important role in both GC initiation and exit [33]. Except for *NFKB2*, most genes encoding for NF-κB signaling components (*NFKB1*, *NFKBIB*, *NFKBID*, *TNFAIP3*) were unaffected by HU or Thy-Noc treatment (Figure 2B). In contrast, expression of *NFKBIA* and *NFKBIE*, two negative regulators of NF-κB signaling, were significantly induced in HU-treated Raji cells (Figure 2B). These indicate that reduced expression of genes encoding the inhibitory components of NF-κB signaling can be important mediator of reduced survival and GCDBL cell fate, aligning with previous reports [33]. Analysis of RNA expression datasets from the AGCBs exhibited higher *NFKBIE* expression within the LZ populations (Figure 2C) [34]. Indeed, the *Nfkbia* and *Nfkbie* expression was higher in mouse splenic plasma B cells which also corresponds to LZs since plasma B cells transit through LZ priors to their differentiation (Figure 2D) [41]. This suggests the inhibition of NF-κB signaling can regulate differentiation of plasma cells transiting through the LZs (Figure 2C, D).

We hypothesized that altered *NFKBIE* status in GC B cells could potentially affect the outcome of the GC reaction. To determine if alterations in NF-κB regulation are associated with patient survival, we compared the overall survival of patients exhibiting inactivating mutations in genes encoding the inhibitory subunits of the NF-κB signaling (*NFKBIB*, *NFKBIE*, and *NFKBID*) using datasets from the cancer genome atlas (TCGA) (Figure 2E, Supplementary Figure 2A, B) [42]. We noted a reduced survival of patients with altered status of *NFKBIB*, *NFKBIE*, and *NFKBID* (Figure 2E, Supplementary Figure 2A, B). Furthermore, analysis of DLBCL patient samples from TCGA datasets revealed frequent mutations in *NFKBIE* (Figure 2F) [42]. These mutations were frequent among diploid DLBCL as 11.5% of diploid DLBCL samples exhibited *NFKBIE* mutation (Figure 2F), aligning with abundance of NF-κB signaling and disease pathogenesis in B-lymphomas[42–44]. The *NFKBIE* mutations found in DLBCL were frameshifts (fs) (*NFKBIE*:Y254Sfs, encoding for 13 amino acids; *NFKBIE*:*L168Pfs* encoding for 42 amino acids), point mutations (*NFKBIE*:*L313Q*, *NFKBIE*:*L441P*, *NFKBIE*:*A251T*), and splicing mutations (X261_splice, A251T) (Figure 2F). These suggest that expression and genetic inactivation of components encoding the inhibitory subunits of NF-κB signaling plays important role in GCDBL fate and DLBCL pathogenesis.

### Inverse correlation between *NFKBIE* expression and *BCL6* in DLBCL pathogenesis

NF**-**κB signaling plays an important role in DLBCL survival and resistance to chemotherapies such as BTK inhibitor, Ibrutinib and R-CHOP (rituximab, cyclophosphamide, doxorubicin, vincristine and prednisone) [30, 45]. To better understand the implication of *NFKBIE* status in DLBLC pathogenesis, we correlated the expression of *NFKBIE* with *BCL6*, a highly amplified and overexpressed gene in DLBCL [15, 46, 47]. Correlation analysis of *BCL6* and *NFKBIE* in DLBCL samples from Gene Expression Profiling Interactive Analysis (GEPIA) indicated a reverse correlation between *BCL6* and *NFKBIE* expression (Pearson correlation R=-0.39) (Figure 3A), suggesting that DLBCLs with higher *BCL6* expression are prone to reduced *NFKBIE* expression (Figure 3A). This suggests that higher NF-κB signaling in B-lymphomas could be associated with frequent inactivation in genes encoding the inhibitory subunits of NF-κB signaling.

**Figure 3:**
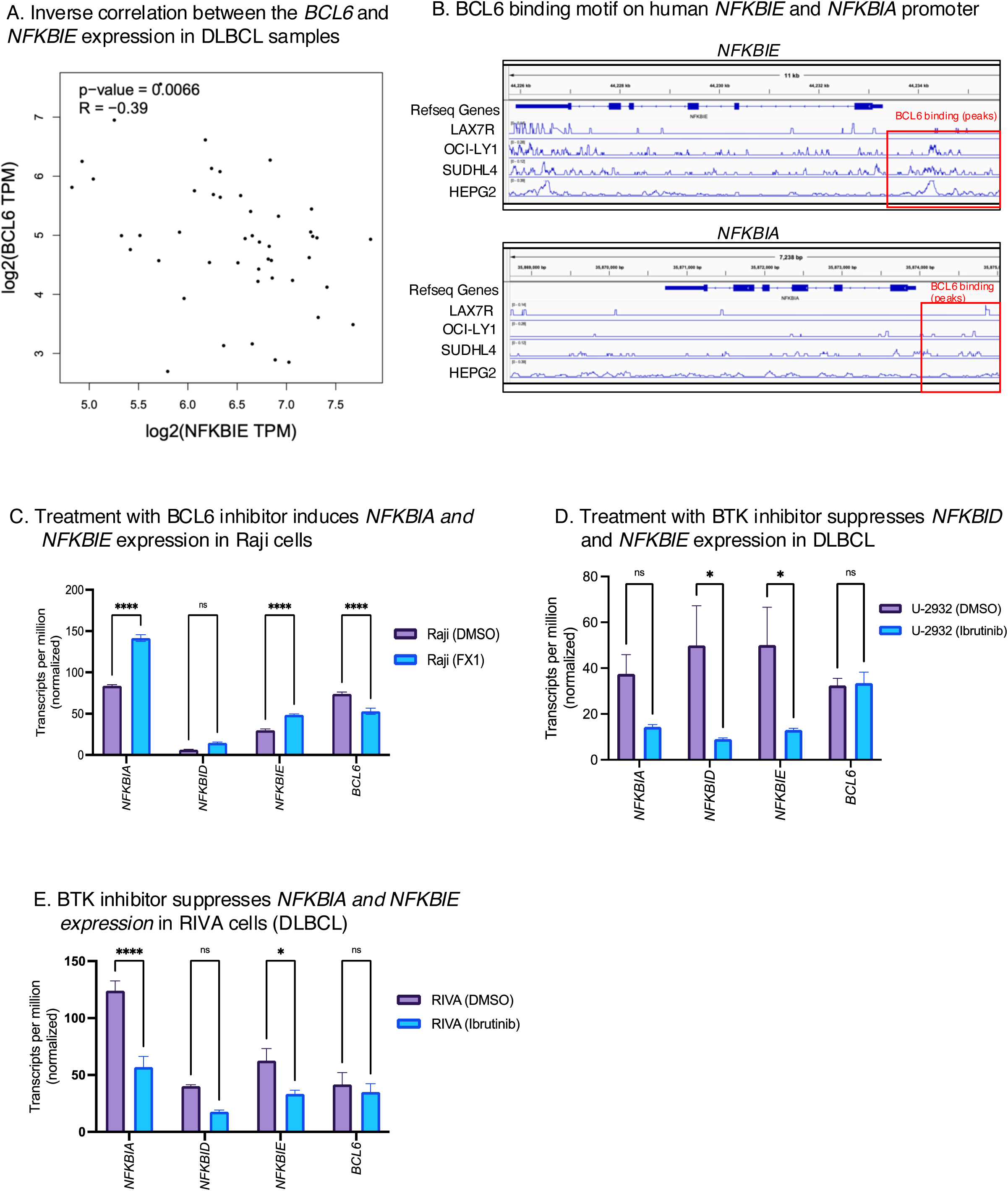
Inverse correlation between *BCL6* and *NFKBIE* in DLBCL. **(A)** Inverse correlation between *BCL6* and *NFKBIE* expression levels in DLBCL samples, analyzed using GEPIA (Pearson correlation: R = −0.39, p = 0.0066). **(B)** BCL6 binding motifs on the human *NFKBIA* and *NFKBIE* promoters were analyzed using ChIP Atlas (https://chip-atlas.dbcls.jp/data/hg38/target/SRX18259603.1.html). BCL6 binding is observed within −2 kilobase of the TSS of *NFKBIA* and *NFKBIE* in LAX7R B-ALL cell (Chip-Atlas#SRX18259603), OCI-LY1 (ChIP-atlas#SRX689470), SU-DHL-4 (ChIP-atlas#SRX4609168) and HepG2 (ChIP-atlas#SRX2636277) cells. BCL6 binding peaks are highlighted within the red rectangle at the promoters of *NFKBIA* and *NFKBIE* in B-ALL, OCI-LY1, SUDHL4, and HepG2 cells. **(C)** FX1 treatment induced *NFKBIA* and *NFKBIE* expression in Raji cells [50]. Normalized TPM values were calculated. Significant differences were observed, with p < 0.0001 for *NFKBIA* and *NFKBIE* when comparing Raji (DMSO) to Raji (FX1). For *NFKBID*, p = 0.0574 in Raji (DMSO) vs. Raji (FX1); Sidak’s multiple comparison test was used (data presented as mean ± SEM, n = 3; GSE254904). **(D)** The BTK inhibitor Ibrutinib suppresses the expression of *NFKBIA*, *NFKBID*, and *NFKBIE* in the U-2932 (DLBCL) cell line [85]. Normalized TPM values were calculated. p-values were 0.2932, 0.0127, 0.0276, and >0.9999 for *NFKBIA*, *NFKBID*, *NFKBIE*, and *BCL6*, respectively, comparing DMSO vs. Ibrutinib-treated U-2932 cells; Sidak’s multiple comparison test was used (data shown as mean ± SEM, n = 6; GSE171763). **(E)** RIVA cells treated with Ibrutinib exhibit reduced expression of *NFKBIA* and *NFKBIE* [85]. P-values of p < 0.0001, 0.1526, 0.0356, and 0.9543 were observed for *NFKBIA*, *NFKBID*, *NFKBIE*, and *BCL6*, respectively, comparing DMSO vs. Ibrutinib-treated groups; Sidak’s multiple comparison test was used (data expressed as mean ± SEM, n = 6; GSE171763).

Since BCL6 binding to *NFKBIE* has been confirmed in GC B cells [48], but whether BCL6 binds to *NFKBIE* in B-lymphomas and other solid cancers is unknown. Loss of BCL6 binding on *NFKBIE* could also lead to induced NF-κB signaling, which is associated with cancer growth and resistance to chemotherapies [49]. We next conducted a BCL6 motif search within the *NFKBIE* promoter using the eukaryotic promoter database and found a highly significant binding of BCL6 at reference position sequence on −830, −883, −913, −916, −950 with cutoff p<0.0001 (data not shown), suggesting a high likelihood of BCL6’s regulatory influence on *NFKBIE* transcription. Furthermore, BCL6 ChIP-sequencing analysis using datasets from Chip-Atlas indicated BCL6 binding on the promoter of *NFKBIA* and *NFKBIE* in LAX7R cells (relapsed B-cell acute lymphoblastic leukemia with *KRAS-G12V* mutation), HepG2 cells (liver hepatocellular carcinoma cell line), as well as in OCI-LY1 and SUDHL4 (DLBCL cells) (Figure 3B). These results suggest that BCL6 binds to the promoter of *NFKBIA* and *NFKBIE* not only in GC B cells but in DLBCL and solid cancers (Figure 3B).

We next asked whether inhibiting the BCL6 activity in B-lymphoma alters the *NFKBIE* expression (Figure 3C). We employed the datasets of Raji cells (Burkitt’s lymphoma) treated with BCL6 inhibitor, FX1 [50] (GSE254904). FX1 binds to lateral groves on BCL6 target DNA sequences and suppresses the repressing complex formation, leading to upregulation of genes suppressed by BCL6 [51]. Raji cells treated with FX1 exhibited increased expression of *NFKBIA* and *NFKBIE* (Figure 3C), suggesting that BCL6 also suppresses *NFKBIA* and *NFKBIE* expression in B-lymphoma (Figure 3C), consistent with its role in suppression of *NFKBIE* in GC B cells [48]. Moreover, the BTK inhibitor Ibrutinib reduced the expression of *NFKBIA*, *NFKBID*, and *NFKBIE* in DLBCL cell lines U-2938, and RIVA cells (Figure 3D-E). These results suggest that chemotherapeutic treatments affect the expression of genes encoding NF-κB inhibitory subunits, which is linked to DLBCL pathogenesis. Inactivating mutations in genes encoding NF-κB inhibitory subunits may serve as an adaptation, promoting higher NF-κB signaling in DLBCL tumors. Conversely, loss of BCL6 regulation of these genes due to chemotherapeutic treatment (BTK inhibitors) may contribute to therapeutic resistance due to enhanced NF-κB signaling. Overall, these findings highlight the role of NF-κB signaling in DLBCL pathogenesis, with BCL6 regulation helping to prevent therapeutic resistance through control of NF-κB signaling.

### Competition between survival and tumor suppressor pathways in HU-treated Raji cells

Multiple signaling pathways--including B cell Receptor (BCR-signaling), NF-κB signaling, p53-response, DNA damage response (DDR) and apoptosis are suppressed by BCL6 in the GC-B cells [15, 29–33, 48]. To examine if transient HU stress in GCDBL affects survival, inflammation, and apoptosis, we mapped these pathways by gene-set enrichment analyses (GSEA) in HU-treated Raji cells (Figure 4A, B, Supplementary Figure 3A, B). We found HU-treatment downregulated gene expression of multiple stress response and survival pathways including DNA damage repair, E2F targets, MYC targets, PI3K/AKT/mTOR, oxidative phosphorylation, hypoxia response, G2-M checkpoint, and mitotic spindle formation (Figure 4A; Supplementary Figure 3A). These results suggest GCDBLs undergoing mild genotoxic stress tend to accumulate in a non-proliferative state (Figure 4A; Supplementary Figure 3A). MYC is required for DZ to LZ entry and GC-exit of B cells [52]. Thus, reduced hallmarks of the MYC pathway in GCDBLs may prevent the GC exit and affinity maturation of GC B cells bearing Burkitt lymphoma signatures. *MYC* is known to be highly expressed in Burkitt lymphomas due to its fusion with the *IG* locus [6, 7]. Suppression of MYC activity in GCDBLs could represent an adaptive mechanism in cases with *MYC* rearrangements, preventing their transition from the DZ to the LZ and, subsequently, inhibiting the GCDBL exit (Figure 4A).

**Figure 4.**
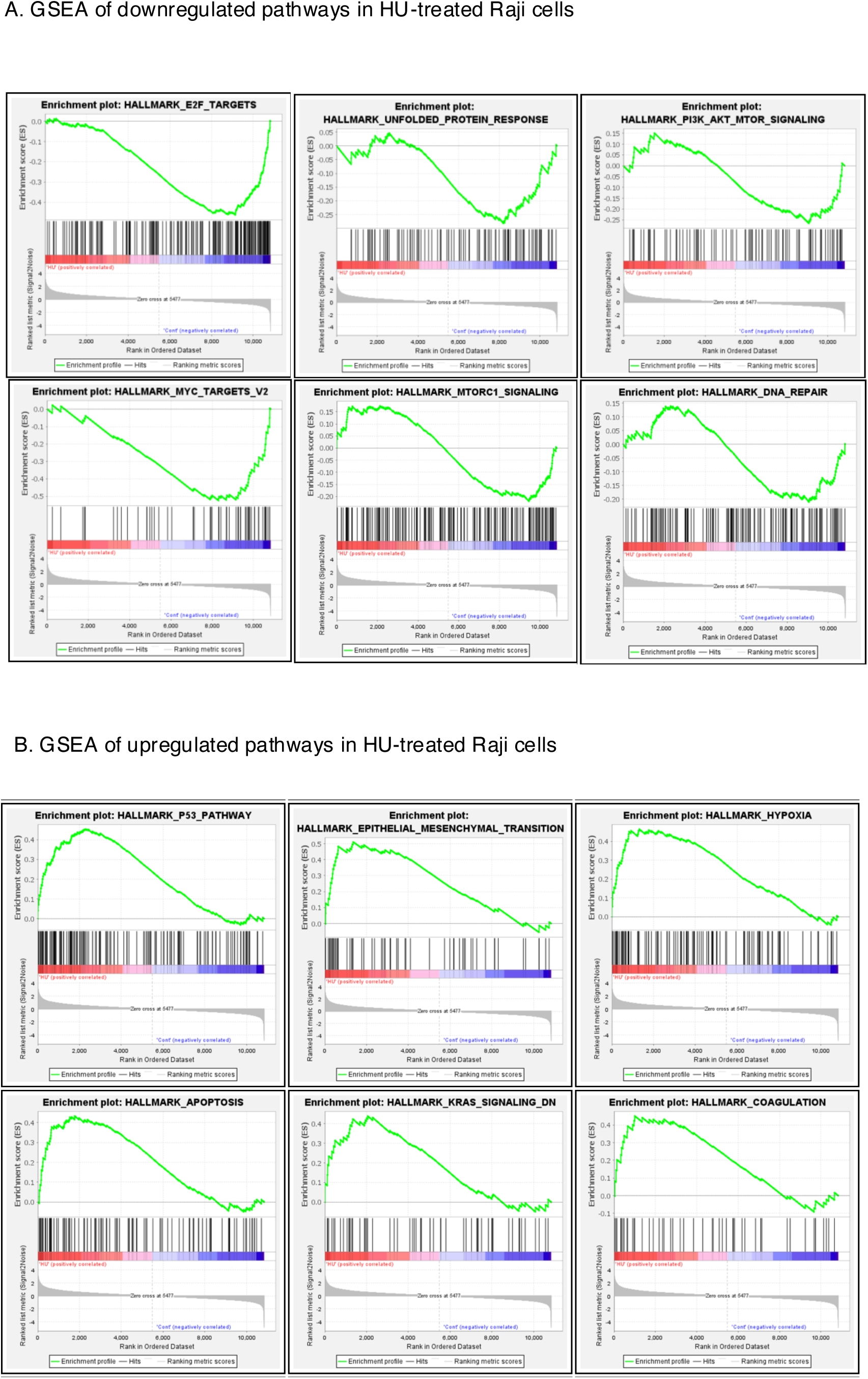
Competition Between Oncogenic and Tumor Suppressor Pathways. **(A)** Gene Set Enrichment Analysis (GSEA) of downregulated pathways in HU-treated Raji cells compared to control cells. The Y-axis represents the enrichment score for pathways, including E2F signaling, unfolded protein response, AKT/mTOR signaling, MYC targets, and DNA repair. Data are the cumulative result of three independent samples per group. **(B)** GSEA of upregulated pathways in HU-treated Raji cells. The Y-axis shows the enrichment score for each pathway, highlighting key pathways such as p53 signaling, epithelial-to-mesenchymal transition, hypoxia, apoptosis, KRAS signaling, and coagulation.

In addition, gene signatures of apoptosis, including the TNF-alpha pathway, were upregulated in HU-treated Raji cells (Figure 4B, Supplementary Figure 3B). Previous reports show that cytokine and inflammatory signaling is suppressed in GC B cells by BCL6 and this suppression is necessary for GC B cell survival in highly apoptotic GC compartments [15, 48]. However, we observed increased gene expression of interferon alpha and gamma-mediated pathway members as well as IL-6, TGF-β, and JAK-STAT3 signaling in HU-treated Raji cells, suggesting the onset of inflammatory signaling in the GCDBLs (Figure 4B, Supplementary Figure 3B). Since BCL6 suppresses these pathways, counteracting BCL6 regulation of inflammatory signaling in GCDBLs could potentially direct them toward an apoptotic fate, facilitating GCDBL elimination.

Somewhat surprisingly, Gene set signatures associated with proliferation pathways, including epithelial to mesenchymal transition (EMT), Apical junction, Notch signaling, and KRAS signaling, were induced in HU-treated Raji cells (Figure 4B, Supplementary Figure 3B). These results suggest that these pathways may actively support the selection and survival of GCDBLs during stress, suggesting their role in GCDBLs resilience and proliferation. The induction of both apoptotic and proliferative pathways in HU-treated Raji cells suggests a competition between apoptotic and survival pathways in GCDBLs. The overall magnitude of apoptotic pathways may dominate survival pathways in the GCDBLs originating in the GCs, leading to their elimination. In contrast, survival pathway signaling could overwhelm pro-apoptotic signals in high affinity B cells which do not harbor oncogenic re-arrangements caused by mis-regulated AID activity in GC B cells undergoing SHM and CSR, allowing high affinity B cells to undergo GC-exit and differentiation during the humoral immune response. Thus, the magnitude of inflammatory, apoptotic, and survival pathways may determine the BCL6 activity in different subpopulations of GC B cells, affecting the GC B cells fate towards apoptosis or differentiation into the plasma and memory B cells.

### *IFNGR1* is a less preferred target of BCL6 compared to its other target gene

BCL6 plays an essential role in promoting the survival of GC B cells and pathogenesis of B-lymphomas by suppressing inflammatory signaling and apoptosis [14, 15]. BCL6 directly binds their promoters with its BTB-domain, then recruits epigenetic silencers thereby repressing transcription of inflammatory signaling genes [48]. Thus, the loss of BCL6 regulation on inflammatory signaling genes in GCDBLs may lead to an apoptotic fate due to the early onset of inflammatory signaling and subsequent activation of apoptosis in the physiological GCs [53, 54]. To evaluate the impact of BCL6 on inflammatory signaling, we analyzed 17 key BCL6 target genes in HU-treated Raji cells [14] (Figure 5A). Despite a trend towards downregulation, *BCL6* levels remained stable in response to HU (Figure 5B), allowing us to identify the genes whose expression is most sensitive to slight modest changes in BCL6 levels (Figure 5A). Among the 17 BCL6 target genes identified (Figure 5A), *IFNGR1*, *IRF7*, and *STAT1* expression were the most induced in HU-treated cells (Figure 5A). This suggesting that *IFNGR1, IRF7* and *STAT1* are particularly sensitive to a modest decline in *BCL6* levels (Figure 5A, B). It is possible that BCL6 association at these promoters is weaker, prompting their swift expression due to mild genotoxic stress (Figure 5A). To confirm these genes are loosely regulated by BCL6, we utilized the datasets of Raji cells treated with FX1, a selective BCL6 inhibitor [50] (Figure 5C, D). Interestingly, *IFNGR1* expression was significantly induced upon FX1 treatment, but *IRF7* and *STAT1* expression were unaltered in FX1-treated Raji cells (Figure 5C). This analysis supports our prediction that BCL6 binding could be easily dissociated on *IFNGR1* in GCDBLs undergoing the chemotherapeutic treatment (Figure 5C).

**Figure 5:**
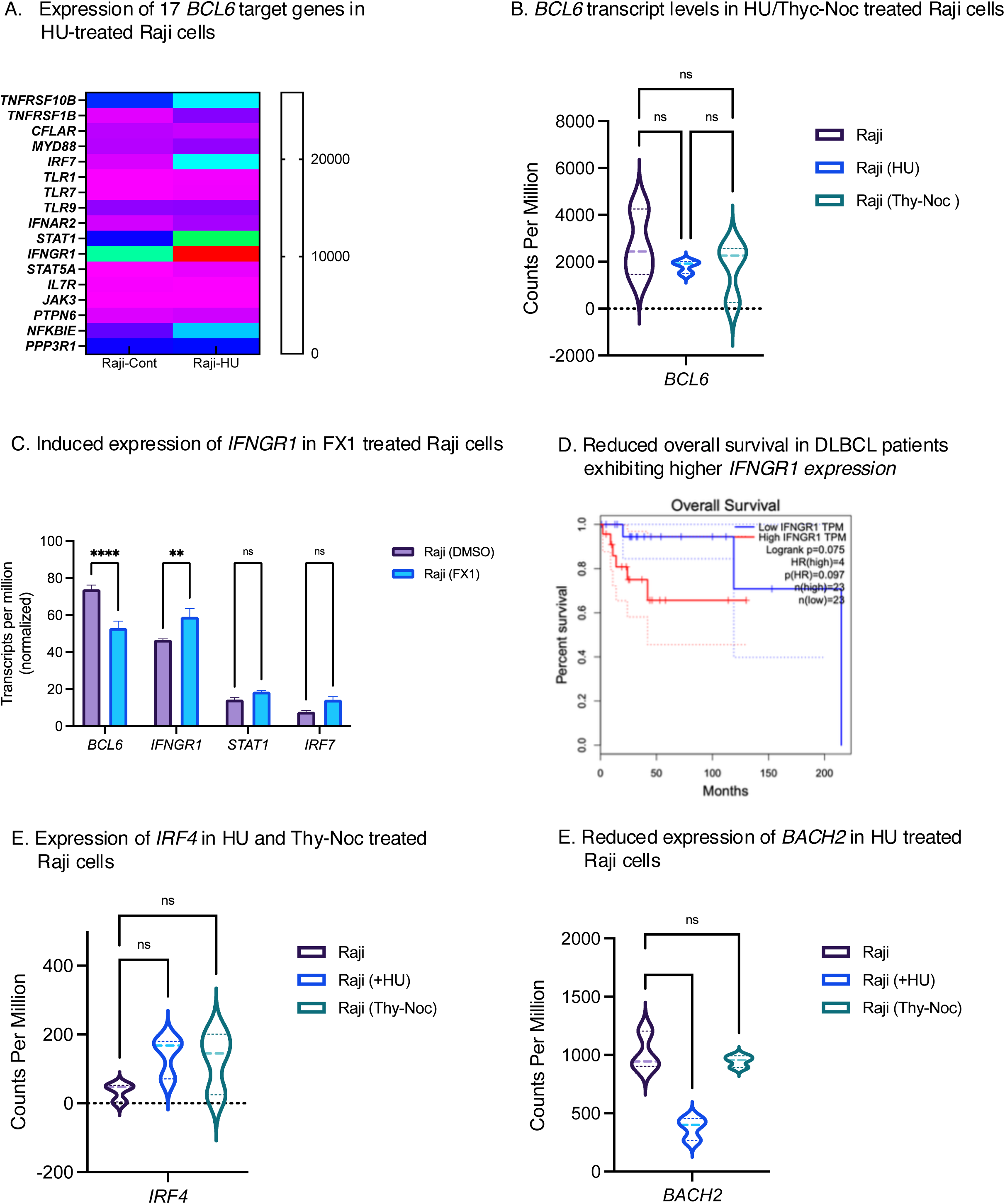
Reduced expression of *IRF7*, *STAT1*, *IFNGR1* and *BACH2* in HU-Treated Raji Cells. **(A)** Expression of BCL6-regulated genes in control and HU-treated Raji cells. Of the 17 key BCL6 target genes analyzed, *IFNGR1*, *STAT1*, and *IRF7* were the most significantly upregulated in response to HU treatment. Statistics: p < 0.0001 for *IFNGR1*, p = 0.0154 for *STAT1*, and p = 0.004 for *IRF7*; P-values were calculated using a two-way ANOVA with multiple comparison testing. Data represent mean ± SEM, n = 3. **(B)** *BCL6* mRNA levels are not significantly altered in Raji (HU) and Raji (Thy-Noc) treated cells compared to control Raji cells: p = 0.7292 for Raji vs. Raji (HU), p = 0.6554 (not significant, ns) for Raji vs. Raji (Thy-Noc), and p = 0.9990 for Raji (HU) vs. Raji (Thy-Noc) using one-way ANOVA with Sidak’s multiple comparison test. Data are presented as mean ± SEM, n = 3. **(C)** BCL6 inhibition further increases *IFNGR1* expression in Raji cells [50]. Normalized TPM values were calculated. Significant differences were observed with p < 0.0001 for *BCL6* when comparing Raji (DMSO) to Raji (FX1). For *IFNGR1*, *STAT1*, and *IRF7*, p = 0.0024, 0.6025, and 0.2127, respectively, in Raji (DMSO) vs. Raji (FX1) using Sidak’s multiple comparison test (data presented as mean ± SEM, n = 3; GSE254904). **(D)** Kaplan-Meier survival analysis comparing *IFNGR1* expression in DLBC groups, defined by the median TPM value (50th percentile). Patients with low *IFNGR1* TPM levels had a trend of higher overall survival compared to those with high *IFNGR1* TPM levels (p-value 0.075). The hazard ratio (HR) for the high *IFNGR1* group was 4 (p = 0.097). n = 23 for both groups. Data are derived from GEPIA database for the DLBC samples **(E)** *IRF4* expression is not altered in Raji (HU) and Raji (Thy-Noc) cells compared to Raji controls. p = 0.1739 (ns) for Raji vs. Raji (HU) and p = 0.2528 (ns) for Raji vs. Raji (Thy-Noc) using one-way ANOVA with Sidak’s multiple comparison test. Data are presented as mean ± SEM, n = 3 **(F)** *BACH2* expression is reduced in Raji (HU) cells but remains unchanged in Raji (Thy-Noc) cells compared to Raji controls. Gene expression is shown as mean CPM values, with statistical significance: p = 0.0008 for Raji vs. Raji (HU) and p = 0.6928 for Raji vs. Raji (Thy-Noc), using one-way ANOVA with Dunnett’s multiple comparison test. Data are presented as mean ± SEM, n = 3.

We found that treatment of Raji cells with BCL6 inhibitors induced the expression of *IFNGR1*, an IFN-γ signaling pathway members (Figure 5C). This suggests a potential mechanism of therapy resistance in clinical cases involving BCL6 and DNA replication inhibitor treatment resulting in loss of BCL6 regulation over *IFNGR1*, which encodes the receptor that drives excess IFN-γ signaling. Induced *IFNGR1* expression can promote IFN-γ signaling, which is associated with DLBCL pathogenesis and decreased efficiency of chimeric antigen receptor (CAR) T-cell therapy [55]. We compared the survival of DLBCL tumors harboring higher *IFNGR1* expression and examined if this was associated with reduced survival than the group exhibiting lower *IFNGR1* expression (Figure 5D). The overall survival was reduced in DLBCL samples exhibiting the higher *IFNGR1* expression, though levels were not significant possibly due to the low number of samples queried (Figure 5D). Similarly, the higher expression of *IRF7* but not *STAT1* exhibited a trend of reduced survival in 23 DLBCL patients exhibiting higher *IRF7* compared to the 23 patients exhibiting lower *IRF7* (Supplementary Figure 4A, B; GEPIA). These results suggest that elevated *IFNGR1* and *IRF7* expression due to loss of BCL6 regulation may contribute to reduced survival in DLBCL patients (Figure 5D, Supplementary Figure 4A).

Given a reciprocal relationship between BCL6 levels with *BACH2*, *IRF4*, and *PRDM1* [56–58], (Sidwell, 2016; 2014; Shinnakasu, 2016), we next examined whether the expression of *BACH2*, *IRF4*, and *PRDM1* was altered in HU and Thy-Noc treated Raji cells (Figure 5E-F, Supplementary Figure 4C,D). Notably, *IRF4* and *PRDM1* expression tend to increase in HU-treated Raji cells, but the increase is not significant (Figure 5E, Supplementary Figure 4C). However, *BACH2* expression was significantly downregulated in HU-treated Raji cells (Figure 5F) indicating a potential blockade in memory B cell differentiation in GCDBLs facing genotoxic stress. This could contribute to the blockade of GCDBL differentiation in B-cell chronic lymphocytic leukemia, a malignancy derived from memory B cells. Moreover, *CREBBP*, a gene encoding for an upstream regulator of BCL6 expression remained unchanged in HU-treated Raji cells (Supplementary Figure 4D), suggesting that BCL6 downregulation via CREBBP downregulation may not be a target mechanism in programmed elimination of GCDBL.

In summary, these results suggest that GCDBLs exposed to transient genotoxic stress may experience a loss of BCL6 regulation preferentially on *IFNGR1*, leading to induced IFN-γ signaling and apoptosis in GCDLB, probing their elimination in the GC microenvironment, while its mis-regulation in DLBCL could promote the disease pathogenesis and therapy resistance due to higher IFN-γ signaling.

### p53 Regulates BCL6 Expression via ATAD2 in B-Lymphomas

The suppression of p53 activity by BCL6 in GC B cells is well-established, promoting the survival of GC B cells [15]. However, BCL6 levels significantly decline as GC B cells exit the GC and differentiate into memory cells and plasmablasts [59–61]. We hypothesized that p53 function may be restored in GC B cells during differentiation, potentially impacting the fate of GC B cells and GCDBLs. To explore this hypothesis, we analyzed the transcriptome of human tonsil GC B cells [34], classifying them into three DZ clusters (DZa, DZb, and DZc), five intermediate zone clusters (INTa, INTb, INTc, INTd, INTe), two LZ clusters (LZa, LZb), a pre-memory group (preM), and two classes of plasmablasts (plasmablast a and b) (Figure 6A). By plotting the log2 fold-change values for *BCL6* and *TP53* transcripts, we found a striking upregulation of *TP53* in LZb cells, suggesting a role for p53 in LZ B cells (Figure 6A). This increase in *TP53* expression coincided with a decline in *BCL6* levels starting from INTd to LZb and from LZb to preM subpopulations (Figure 6A). The inverse relationship between *BCL6* and *TP53* levels aligns with the role of BCL6 in p53 suppression [15]. However, our observation of induced *TP53* in LZb subpopulation provides additional insight on the emerged regulation of p53 could be specific to GC B undergoing selection and terminal differentiation (Figure 6A). This suggests that *TP53* is upregulated in LZ B cells before their differentiation into memory and plasma cells (Figure 6A).

**Figure 6.**
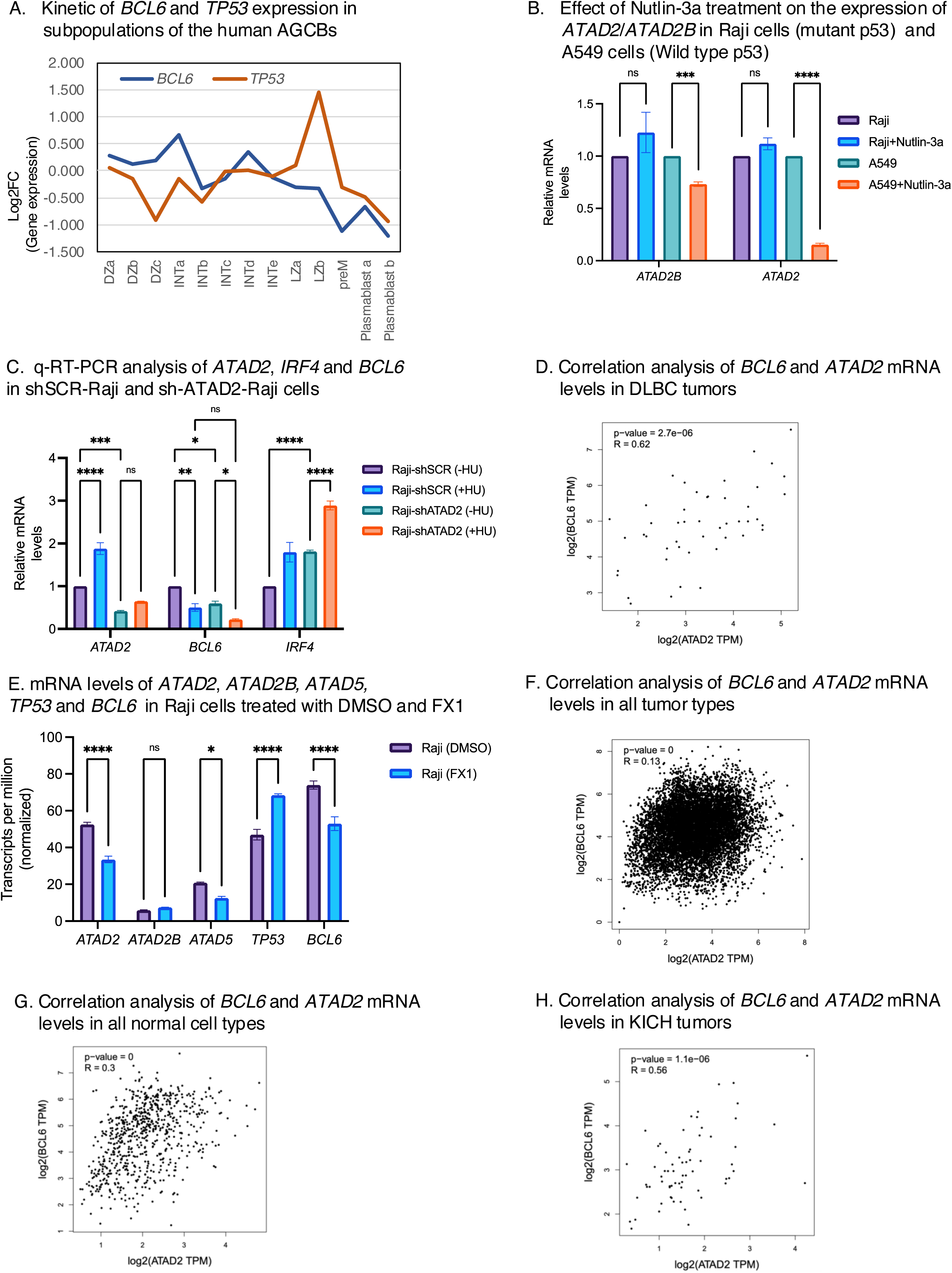
p53 suppresses *ATAD2* expression in GCDBL. **(A)** Kinetics of *BCL6* and *TP53* mRNA expression in a subpopulation of human tonsil AGCBs. *TP53* expression peaks in the LZb subpopulation, while *BCL6* expression is reduced at the same point **(B)** Effects of Nutlin-3a treatment in Raji and A549 cells and quantitation of *ATAD2* and *ATAD2B* expression levels by real time quantitative polymerase chain reaction (RT-qPCR). Raji and A549 cells were treated with vehicle or Nutlin-3a (10μM) for 24 hours followed by qRT-PCR analysis. For *ATAD2B*, p= 0.3076 (ns) for Raji vs Raji (Nutlin-3a) and p= 0.0003 for A549 vs A549 (Nutlin-3a) treated groups. For *ATAD2* p= 0.1079 (ns) for Raji vs Raji (Nutlin-3a) and p<0.0001 for A549 vs A549 (Nutlin-3a) treated groups using unpaired t-test. Data are presented as mean ± SEM, n=3 **(C)** Analysis of *ATAD2*, *BCL6*, and *IRF4* expression in shSCR-Raji and shATAD2-Raji cells. shSCR-Raji and shATAD2-Raji cells were treated with 10 mM HU for 24 hrs. mRNA levels of *ATAD2*, *IRF4* and *BCL6* were measured by RT-qPCR. Tukey’s multiple comparison test. For *ATAD2*; p=0.0006 for shSCR-Raji (-HU) vs shATAD2-Raji cells (-HU). p<0.0001 for shSCR-Raji (-HU) vs shSCR-Raji (+HU). p=0.2869 for shATAD2-Raji (-HU) vs shATAD2-Raji (+HU). For *BCL6*, p= p=0.0029 for shSCR-Raji (-HU) vs shSCR-Raji cells (+HU). p=0.0185 for shSCR-Raji (-HU) vs shATAD2-Raji (-HU) cells. p=0.03 for shATAD2-Raji (-HU) vs shATAD2-Raji (+HU) cells. For *IRF4*, p<0.0001 for shSCR-Raji (-HU) vs shSCR-Raji cells (+HU). p<0.0001 for shSCR-Raji (-HU) vs shATAD2-Raji (-HU). p<0.0001 for shATAD2-Raji (-HU) vs shATAD2-Raji (+HU). Tukey’s multiple comparison test. Data are presented as mean ± SEM, n=3. **(D)** Correlation between *ATAD2* and *BCL6* expression in DLBC tumors. Scatter plot showing the correlation between *ATAD2* and *BCL6* gene expression levels in DLBC tumor samples, generated using GEPIA. The Pearson correlation coefficient (R = 0.62) indicates a positive correlation between the two genes. p=2.7e−06 **(E)** Normalized transcript levels of *ATAD2*, *ATAD2B, ATAD5, TP53* and *BCL6* in Raji cells treated with DMSO or FX1 [50] (GSE254904). *ATAD2*; p<0.0001 for Raji (DMSO) vs Raji (FX1). *ATAD2B*; p=0.9808 for Raji (DMSO) vs Raji (FX1). *ATAD5*; p=0.0115 for Raji (DMSO) vs Raji (FX1). *TP53*; p<0.0001 for Raji (DMSO) vs Raji (FX1). *BCL6*; p<0.0001 for Raji (DMSO) vs Raji (FX1). Sidak’s multiple comparison test was used. Normalized TPM values were calculated (data expressed as mean ± SEM, n = 3; GSE254904). **(F)** Correlation between *ATAD2* and *BCL6* expression across all tumor types analyzed. Scatter plot showing the correlation between *ATAD2* and *BCL6* gene expression all tumor types analyzed using GEPIA (R = 0.13, p = 0). **(G)** Correlation between *ATAD2* and *BCL6* expression across normal cell types analyzed. Scatter plot showing the correlation between *ATAD2* and *BCL6* gene expression in normal cell types using GEPIA (R = 0.30, p = 0). **(H)** Correlation between *ATAD2* and *BCL6* expression in Kidney Chromophobe (KICH) samples, using GEPIA database (R = 0.56, p = 1.1e−06).

To investigate the functional significance of p53 upregulation in LZb cells, we identified p53-dependent genes among the differentially expressed genes (DEGs) in HU-treated Raji cells (Supplementary Figure 5A, B). We found 81 p53-dependent DEGs in HU-treated cells and 9 in thymidine/nocodazole (Thy-Noc)-treated cells (Supplementary Figure 5A, B). Notably, *ATAD2B*, a bromodomain protein, was induced in HU-treated cells (Supplementary Figure 5A), suggesting that p53 fails to suppress *ATAD2B* under these conditions (Supplementary Figure 6A). Given the presence of *TP53* mutations (*TP53R213Q* and *TP53Y234H*) in Raji cells, we hypothesized that mutant p53 is unable to suppress *ATAD2B* in HU-treated Raji cells, resulting in its upregulation. To test if wild-type p53 can suppress *ATAD2B*, we treated A549 (wild-type p53) cells with Nutlin-3a, a p53 agonist that stabilizes p53 by inhibiting its interaction with MDM2 [62]. Nutlin-3a had no effect on *ATAD2B* levels in Raji cells but significantly reduced its expression in A549 cells (Figure 6B). Interestingly, *ATAD2*, another member of similar family, was significantly suppressed in Nutlin-3a-treated A549 cells but not in Raji cells, indicating that wild-type p53 suppresses *ATAD2* expression (Figure 6B). These results suggest that p53 downregulates *ATAD2* expression in GC B cells, whereas mutant p53 in Raji cells lacks this function, leading to dysregulation of *ATAD2* expression (Figure 6A, B). *ATAD2* is an oncogene upregulated in multiple cancers and interacts with oncogenic transcription factors associated with B-lymphomagenesis [63]. This suggests a tumor suppressor role for p53 through the suppression of ATAD2.

As *TP53* expression is induced in activated GC B cells transitioning within LZs (Figure 6A), we hypothesized that p53 may play a role in GC B cells transitioning from the LZ to pre-memory and plasmablast stages. With the concurrent reduction in *BCL6* expression at this stage (Figure 6A) [64], we hypothesized that p53 suppresses *BCL6* expression in GC B cells via an ATAD2-dependent manner to ensure their quality control prior to GC exit and terminal differentiation into memory and plasma cells. By removing the *BCL6* barrier, p53-mediated ATAD2 suppression may enable the exit of GC B cells harboring high-affinity BCRs. To test this, we performed *ATAD2* knockdown in Raji cells using short-hairpin RNA (shATAD2-Raji) and compared *BCL6* levels to those in scramble-RNA knockdown Raji cells (shSCR-Raji) (Figure 6C). shATAD2-Raji cells showed significantly reduced *ATAD2* expression (Figure 6C). Importantly, *BCL6* expression was also reduced in shATAD2-Raji cells, especially upon HU treatment, suggesting that *ATAD2* is necessary for full *BCL6* expression during genotoxic stress (Figure 6C). This reduction in *BCL6* correlated with an increase in *IRF4* levels (Figure 6C) and is consistent with the inverse relationship between *IRF4* and *BCL6* [65, 66]. These findings imply that p53 suppresses *BCL6* expression by inhibiting *ATAD2* transcription.

Given the elevated *BCL6* levels and frequent p53 mutations in Raji cells as well as multiple tumors [67, 68], we next examined the correlation between *BCL6* and *ATAD2* expression in DLBCL samples (Figure 6D). Data from the GEPIA database showed a significant positive correlation between *ATAD2* and *BCL6* expression in DLBCL samples (Pearson R = 0.62, p = 2.7e−06) (Figure 6D). Indeed, Raji cells treated with the BCL6 inhibitor FX1 exhibited reduced *ATAD2* and *ATAD5* expression, suggesting that BCL6 is required for their expression [15, 50] (Figure 6E). FX1 treatment of Raji cells confirmed a reduction of *BCL6* and an increase in *TP53* expression, consistent with BCL6’s known role in p53 suppression (Figure 6E) [15]. In contrast, a weaker correlation between *ATAD2* and *BCL6* was observed across all tumor types analyzed (R = 0.13, p = 0) and in normal samples (R = 0.30, p = 0) (Figure 6F, G), suggesting that the *ATAD2*-*BCL6* correlation may be specific to DLBCL (Figure 6D-G). A similar correlation was also observed in kidney chromophobe (KICH) and cervical squamous cell carcinoma samples, with R-values of 0.46 (p = 0) and 0.56 (p = 1.1e−06), respectively (Figure 6H, Supplementary Figure 5C), indicating a positive correlation between *ATAD2* and *BCL6* expression in specific solid tumors as well as B-lymphomas.

We observed a positive correlation between *BCL6* and *ATAD2* expression in tumors (Figure 6D, F, G, H). Since *Bcl6* expression is reduced in plasma B cells compared to the GC B cells, we hypothesized that expression of *ATAD* family members could also be reduced in plasma B cells. We examined this hypothesis using human and mice GC B cell dataset [34, 41]. We confirmed that *BCL6* expression was lowest in plasma cells compared to GC B cells in both human and mouse datasets analyzed (Supplementary Figure 5D, E). We observed a trend of reduced *ATAD2B* expression but not *ATAD2* expression in human GC B cells, when comparing the LZb and plasmablast B population (Figure 5D). On the other hand, *Atad2*, *Atad2b*, and *Atad5* expression was reduced in mouse plasma cells, correlating well with the decline in *Bcl6* (Supplementary Figure 5D). These observations further suggest coordinated expression of ATAD family members and BCL6 in GC B cells.

### Expression of genes encoding for GPCRs, G-proteins and *RAC1*/*RHO* proteins in HU treated Burkitt’s lymphoma Cells

We next examined whether GPCRs and associated receptors were altered in HU- and Thy-Noc-treated Raji cells, since GPCR signaling remains essential for B cell migration and survival [69–71]. Expression of most GPCRs and other receptors including *CXCR4*, CXCR5, *CXXC1*, *CXXC5*, *S1PR2*, *P2RY8*, *CD38*, *CD80*, *CD82*, *CD83*, *CD84*, *CD86*, and *CD99* showed stable expression regardless of treatments (Figure 7A). *CCR7* and *CD81* expression increased following HU stress, suggesting their activity may play critical roles in GCDBL survival when facing genotoxic stress in GC microenvironment. We further analyzed G-protein subunit expression across three classes, Gα, G_β_ and Gγ (Figure 7B-D). Among Gα proteins, *GNAI2* was significantly upregulated in HU-treated cells, while the expression of *GNAS*, *GNA11*, *GNA12*, *GNAI3*, and *GNAZ* remained unchanged (Figure 7B). Among Gβ subunits, only *GNB2* expression was significantly reduced in HU and Thy-Noc-treated Raji cells, whereas *GNB1*, *GNB3*, *GNB4*, and *GNB5* were unaffected by either treatment (Figure 7C). No significant changes were observed for Gγ subunits *GNG2*, *GNG5*, *GNG7*, *GNG10*, and *GNGT2* in response to HU or Thy-Noc treatment (Figure 7D). These results suggest that GPCR expression and their associated G-protein subunits are generally unaffected by mild genotoxic stress or Thy-Noc treatment, indicating that they may not be regulated in a cell-cycle-specific manner in GCDBLs (Figure 7A-D). The increased surface expression of *CD81* and *CCR7* upon HU treatment (Figure 7A) supports the idea that GPCR signaling remains constitutively active in GCDBLs (Figure 7A). In addition, no substantial downregulation of GPCRs was observed, suggesting that GPCR expression and possibly GPCR signaling is not compromised in the GCDBLs undergoing genotoxic stress and elimination.

**Figure 7.**
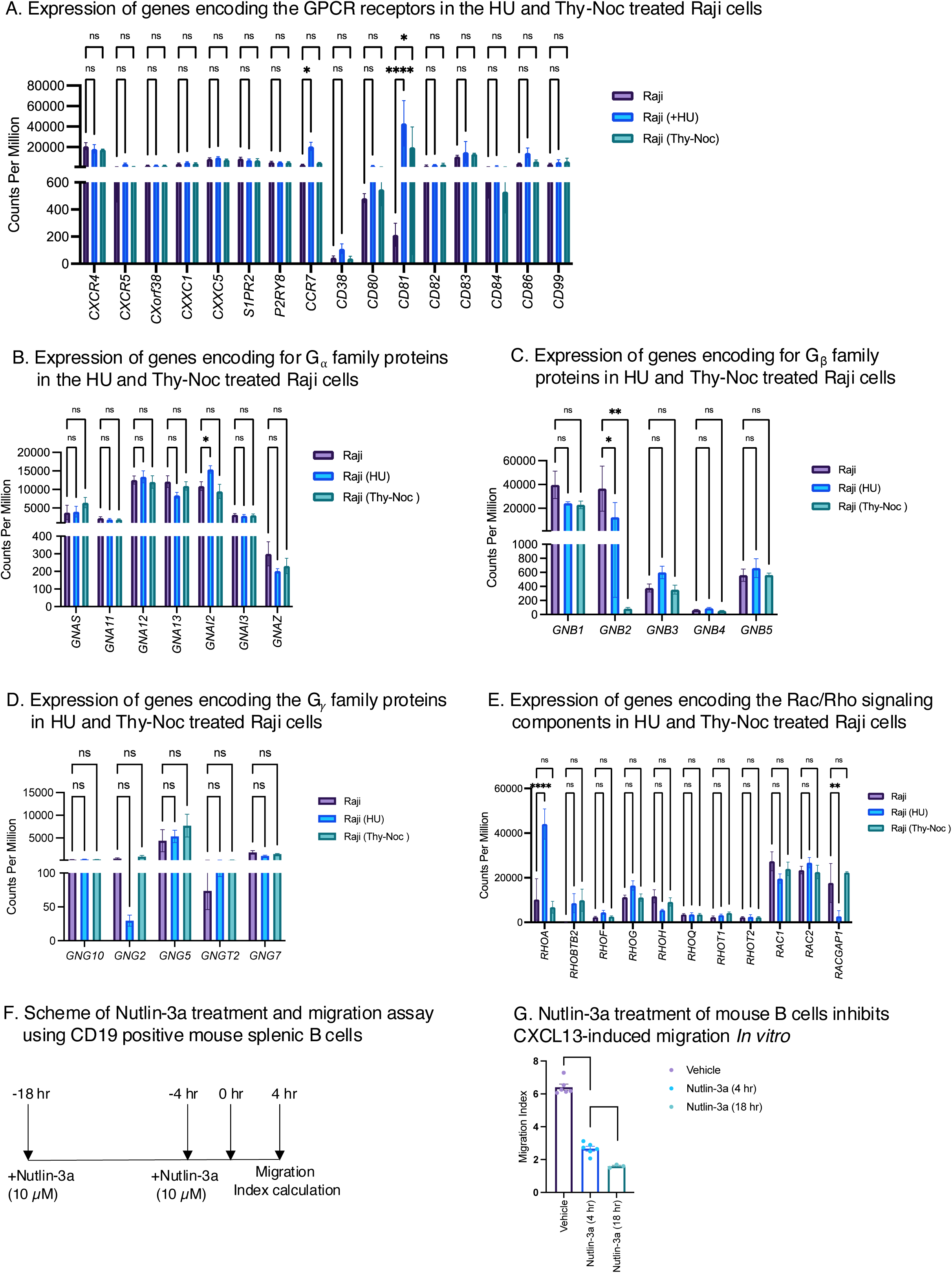
Expression of genes encoding the GPCR and G-protein in the Raji (HU) and Raji (Thy-Noc) treated cells. **(A)** Transcript levels for *CXCR4*, *CXCR5*, *CXorf38*, *CXXC1*, *CXXC5*, *S1PR2*, *P2RY8*, *CCR7*, *CD38*, *CD80*, *CD81*, *CD82*, *CD83*, *CD84*, *CD86*, *CD99* in Raji, Raji (HU) and Raji (Thy-Noc) cells were measured from TPM values. *CCR7* and *CD81* were significantly induced in Raji vs Raji (HU) cells. for *CCR7*; p=0.0272 for Raji vs Raji (HU) and p= 0.9587 for Raji vs Raji (Thy-Noc). For *CD81*; p<0.0001 for Raji vs Raji (HU) and p= 0.0123 for Raji vs Raji (Thy-Noc). For *CXCR4*; p=0.9179 for Raji vs Raji (HU) and p= 0.8882 for Raji vs Raji (Thy-Noc) For *CXCR5*; p=0.8880 for Raji vs Raji (HU) and p= 0.9986 for Raji vs Raji (Thy-Noc). For *CXorf38*; p=0.9992 for Raji vs Raji (HU) and p>0.9999 for Raji vs Raji (Thy-Noc). For *CXXC1*; p=0.9783 for Raji vs Raji (HU) and p= 0.9986 for Raji vs Raji (Thy-Noc). For *CXXC5*; p=0.9767 for Raji vs Raji (HU) and p= 0.9948 for Raji vs Raji (Thy-Noc). For *S1PR2*; p=0.9740 for Raji vs Raji (HU) and p= 0.9647 for Raji vs Raji (Thy-Noc). For *P2RY8*; p>0.9999 for Raji vs Raji (HU) and p= 0.9995 for Raji vs Raji (Thy-Noc). For *CD38*; p>0.9999 for Raji vs Raji (HU) and p>0.9999 for Raji vs Raji (Thy-Noc). For *CD80*; p=0.9783 for Raji vs Raji (HU) and p>0.9999 for Raji vs Raji (Thy-Noc). For *CD82*; p=0.9798 for Raji vs Raji (HU) and p= 0.9707 for Raji vs Raji (Thy-Noc). For *CD83*; p=0.8099 for Raji vs Raji (HU) and p= 0.9197 for Raji vs Raji (Thy-Noc). For *CD84*; p=0.9946 for Raji vs Raji (HU) and p= 0.9999 for Raji vs Raji (Thy-Noc). For *CD86*; p=0.3016 for Raji vs Raji (HU) and p= 0.9670 for Raji vs Raji (Thy-Noc). For *CD99*; p=0.9639 for Raji vs Raji (HU) and p= 0.9313 for Raji vs Raji (Thy-Noc) using 2-way ANOVA with Tukey’s multiple comparison test. Data are presented as mean ± SEM, n=3. **(B)** Transcript levels of G_α_ subunits in Raji, Raji (HU) and Raji (Thy-Noc) cells were measured from TPM values. Expression of other G_α_ subunits *GNAS*, *GNA11*, *GNA12*, *GNA13*, *GNAI3*, *GNAZ* was unaltered between Raji, Raji (HU) and Raji (Thy-Noc) cells. For *GNAS*; p=0.9874 for Raji vs Raji (HU) and p= 0.2161 for Raji vs Raji (Thy-Noc). For *GNA11*; p=0.9751 for Raji vs Raji (HU) and p= 0.9720 for Raji vs Raji (Thy-Noc). For *GNA12*; p=0.8411 for Raji vs Raji (HU) and p= 0.9429 for Raji vs Raji (Thy-Noc). For *GNA13*; p=0.0588 for Raji vs Raji (HU) and p= 0.7378 for Raji vs Raji (Thy-Noc). For *GNAI2*; p=0.0171 for Raji vs Raji (HU) and p= 0.6431 for Raji vs Raji (Thy-Noc). For *GNA13*; p=0.9835 for Raji vs Raji (HU) and p= 0.9969 for Raji vs Raji (Thy-Noc). For *GNAZ*; p=0.9979 for Raji vs Raji (HU) and p= 0.9989 for Raji vs Raji (Thy-Noc). 2way ANOVA. Tukey’s multiple comparison test. Data are presented as mean ± SEM, n=3. **(C)** Transcript levels of G_β_ family subunits in Raji, Raji (HU) and Raji (Thy-Noc). For *GNB1*; p=0.2417 for Raji vs Raji (HU), p= 0.1890 for Raji vs Raji (Thy-Noc). For *GNB2*; p=0.0396 for Raji vs Raji (HU), p= 0.0014 for Raji vs Raji (Thy-Noc). For *GNB3*; p=0.9997 for Raji vs Raji (HU), p>0.9999 for Raji vs Raji (Thy-Noc). For *GNB4*; p>0.9999 for Raji vs Raji (HU), p= 0>0.9999 for Raji vs Raji (Thy-Noc). For *GNB5*; p>0.9999 for Raji vs Raji (HU), p>0.9999 for Raji vs Raji (Thy-Noc). 2way ANOVA. Tukey’s multiple comparison test. Data are presented as mean ± SEM, n=3. **(D)** Transcript levels of G_ψ_ family subunits in Raji, Raji (HU) and Raji (Thy-Noc) cells. mRNA expression of G_ψ_ subunits *GNG10*, *GNG2*, *GNG5*, *GNGT2*, and *GNG7* was analyzed using the TPM values in Raji, Raji (HU) and Raji (Thy-Noc) groups. For *GNG10*; p=0.9990 for Raji vs Raji (HU), p=0.9997 for Raji vs Raji (Thy-Noc). For *GNG2*; p=0.9452 for Raji vs Raji (HU), p=0.9499 for Raji vs Raji (Thy-Noc). For *GNG5*; p=0.7741 for Raji vs Raji (HU), p=0.0578 for Raji vs Raji (Thy-Noc). For *GNGT2*; p=0.9994 for Raji vs Raji (HU), p=0.9985 for Raji vs Raji (Thy-Noc). For *GNG7*; p=0.8307 for Raji vs Raji (HU), p=0.9657 for Raji vs Raji (Thy-Noc). 2way ANOVA. Tukey’s multiple comparison test. Data are presented as mean ± SEM, n=3. **(E)** transcript levels of *RHOA*, *RHOBTB2*, *RHOF*, *RHOG*, *RHOH*, *RHOQ*, *RHOT1*, *RHOT2*, *RAC1*, *RAC2* and *RACGAP1* in Raji, Raji (HU) and Raji (Thy-Noc) cells. *RHOA*; p<0.0001 for Raji vs Raji (HU) and p= 0.7265 for Raji vs Raji (Thy-Noc). *RHOBTB2*; p=0.1545 for Raji vs Raji (HU) and p= 0.0840 for Raji vs Raji (Thy-Noc). *RHOF*; p=0.8819 for Raji vs Raji (HU) and p= 0.9981 for Raji vs Raji (Thy-Noc). *RHOG*; p=0.4877 for Raji vs Raji (HU) and p= 0.9990 for Raji vs Raji (Thy-Noc). *RHOH*; p=0.3647 for Raji vs Raji (HU) and p= 0.8411 for Raji vs Raji (Thy-Noc). *RHOQ*; p=0.9998 for Raji vs Raji (HU) and p> 0.9999 for Raji vs Raji (Thy-Noc). *RHOT1*; p=0.9781 for Raji vs Raji (HU) and p= 0.9102 for Raji vs Raji (Thy-Noc). *RHOT2*; p=0.9988 for Raji vs Raji (HU) and p> 0.9999 for Raji vs Raji (Thy-Noc). *RAC1*; p=0.2071 for Raji vs Raji (HU) and p= 0.7322 for Raji vs Raji (Thy-Noc). *RAC2*; p=0.7467 for Raji vs Raji (HU) and p= 0.9805 for Raji vs Raji (Thy-Noc). *RACGAP1*; p=0.0045 for Raji vs Raji (HU) and p= 0.5554 for Raji vs Raji (Thy-Noc). 2way ANOVA. Tukey’s multiple comparison test. Data are presented as mean ± SEM, n=3. **(F)** Schematic representation of CD19-positive mouse B cells treatment with Nutlin-3a (10 μM) for 4 and 18 hours. (G) Assessment of migration against the rmCXCL13 ligand (1000 ng/μl) after 4 and 18 hours of Nutlin-3a treatment. The migration index represents the normalized value of cells migrating in transwells containing rmCXCL13 compared to control medium wells without the ligand. Nutlin-3a (10 μM) treatment for 4 hours reduced the migration index, with a further reduction observed after 18 hours. Data are presented as mean ± SEM, n = 6 for vehicle and Nutlin-3a (4 hours) groups. n=3 for Nutlin-3a (18 hours). p<0.0001 for Vehicle vs Nutlin-3a (4 hours) and p=0.0019 for Nutlin-3a (4 hours) vs Nutlin-3a (18 hours). Unpaired t-test.

GPCR signaling assists B cell migration while Rho family proteins negatively regulate cell migration [72–74], therefore we next examined their transcriptional regulation. Expression of *RHOA,* a p53 dependent inhibitor of cell migration, was significantly induced in HU-treated Raji cells (Figure 7E), indicating that mild genotoxic stress may negatively regulate B-cell migration in GCDBLs. Furthermore, expression of *RACGAP1*, whose product promotes migration, was reduced [73] (Figure 7E), suggesting a reduced tendency for migration in GCDBLs undergoing genotoxic stress. These findings indicate that migration of GCDBLs could be affected in GCDBL experiencing genotoxic stress, potentially limiting lymphoma cell migration and hindering GC exit. Given the induced expression of the negative regulator *RHOA*, and reduced expression of the positive regulator *RACGAP1* (Figure 7E), we hypothesized that p53 may regulate the GC B cell migration. We sorted CD19-positive mouse splenic B cells and tested their migration towards recombinant mouse CXCL13 (rmCXCL13) ligand *in vitro* (Figure 7F, G). Mouse B cells showed a higher migration index towards the rmCXCL13, however Nutlin-3a treatment impaired CXCL13 ligand-dependent migration (Figure 7G). Nutlin-3a induced impairment of CXCL13 migration was time dependent, as migration index of 4 hours of Nutlin-3a treated B cell was higher than 18 hours treated B cells (Figure 7G). Together, these results suggest that p53 restricts B cell migration towards the CXCL13 ligand *i.e.* LZs, implying a critical role of p53 in LZ B cells and GC B cell undergoing terminal differentiation.

## Discussion

In this study, by inducing mild genotoxic stress with HU in Raji cells, we investigated the pathways governing GC B cell fate determination for cells harboring oncogenic rearrangements such as *IG-MYC* and *IG-BCL6*. The HU stress, inducing a GC-like stress similar to that experienced by B cells undergoing CSR and SHM, allowed us to map the transcriptional regulation of key signaling pathways, including B cell receptor (BCR), NF-κB, BCL6 and p53 target genes, as well as GPCRs and G proteins (Figure 1A,B). Our comparative analyses with gene expression profiles from normal human tonsillar GC B cells and DLBCL patient samples provided novel insights highlighting the altered dynamics and genetic mutations associated with NF-κB activation, IFN-γ signaling, BCL6, and p53 inactivation in the GCDBL fate and DLBCL pathogenesis.

### BCR Signaling and NF-κB signaling in HU-Treated Raji Cells

Our findings indicate that the expression of genes encoding the BCR signaling components remain largely unchanged in HU-treated Raji cells (Figure 2A). This suggests that BCR signaling is constitutively active in both GC B cells and GCDBLs, indicating that it is not a dominant regulator of GCDBL elimination in GC microenvironment. While we noted no significant changes in the expression of NF-κB signaling components, we observed an upregulation of its inhibitory subunits, *NFKBIA* and *NFKBIE* (Figure 2B). This suggests that genotoxic stress inhibits NF-κB signaling, subsequently influencing the fate of GCDBLs, although this needs further study in mouse models to determine if enhanced NF-κB reduces survival for GC B cells residing in DZ and LZ compartments. The inverse correlation between *BCL6* and *NFKBIE* expression in DLBCL samples (GEPIA) suggests that BCL6 may promote NF-κB signaling in clinical DLBCLs (Figure 3A). Additionally, Burkitt’s lymphoma and DLBCL lines treated with BCL6 inhibitors, BTK inhibitors and R-CHOP inhibitors showed reduced expression of genes encoding NF-κB inhibitory subunits (Figure 3D-F), indicating a possible mode of therapeutic resistance caused by BTK inhibitors resulting from increased NF-κB signaling [30, 45, 75]. Furthermore, we found that 11.3% of diploid DLBCL cancers and 10.8% of total DLBCL cancers (TCGA) exhibited *NFKBIE* mutations (Figure 2F), indicating that inactivating mutations in NF-κB inhibitory subunits represent a potential adaptive mechanism in DLBCL tumorigenesis due to higher NF-κB singling. These findings highlight NF-κB inhibition as a promising strategy for treating GCDBLs in combination with therapies such as R-CHOP, BTK inhibitors, and BCL6 inhibitors.

### Loss of BCL6 Regulation on *IFNGR1*

We observed increase in *IFNGR1* expression in B-lymphomas treated with HU, BCL6 inhibitor, and BTK inhibitors, suggesting these drugs elevate IFN-γ signaling and contribute to therapeutic resistance, as IFN-γ signaling is implicated in chemotherapeutic resistance and PD1-blockade [76, 77]. Our comparative analysis reveals the early onset of NF-κB and IFN-γ signaling in GCDBLs due to loss of BCL6 regulation, which could suppress GCDBL survival within the GC microenvironment; however, hijacked regulation evidenced by *NFKBIE* mutations and easy dissociation of BCL6 binding on the *IFNGR1* indicates that these serve as adaptations in B-lymphoma survival and therapeutic resistance (Figure 2D-F, 5C-D). These adaptation mechanisms could be fundamental for B-lymphoma survival, influencing their elimination within the GC microenvironment and facilitating their escape from the GC barrier. Increased IFN-γ signaling in GC B cells within the GC microenvironment may also promote autoimmune responses in B cells [78, 79]. Additionally, the loss of BCL6 regulation on these inflammation-related genes may drive pathogenesis and therapy resistance in B-lymphomas, potentially serving as an adaptive mechanism for tumor proliferation.

### New Roles of p53 in GC B Cell Fate Determination

A key observation in our study was the upregulation of p53 expression coinciding with the differentiation of GC B cells and the downregulation of BCL6 in human and mouse GC B cells (Figure 6A, Supplementary Figure 5F) [34, 41]. This finding supports a novel role for p53 in GC B cell differentiation, suggesting that restoration of p53 function is essential to ensure quality control in GC B cells undergoing terminal differentiation. This regulation may prevent affinity maturation of GCDBLs and B-lymphomagenesis, aligning with p53’s role in suppressing spontaneous lymphoma generation [80]. Future studies with conditional depletion of *Trp53* in LZ B cells in mouse models can further define this role. Moreover, our findings indicate that p53 may indirectly suppress BCL6 expression via ATAD2 downregulation in GC B cells undergoing terminal differentiation (Figure 6A, 6C, Supplementary Figure 5D-E). This suppression may limit GCDBL survival due to reduced BCL6 levels, thereby reducing the persistence of DLBCL and follicular lymphoma precursors prior to GC exit. As p53 suppresses *ATAD2* expression (Figure 6B), this function may also reduce the frequency of AID-induced genomic rearrangements in GC B cells, given the role of bromodomain proteins in facilitating translocations during the CSR-induced DNA breaks [28]. Thus, ATAD2 may play additional roles in GC B cells related to mutagenic DNA repair [81]. Indeed, ATAD2 promotes oncogenic transcription by collaborating with MYC, E2F1, E3F3 [63, 81, 82]. ATAD2 binds acetylated histone H3K14 motifs through its bromodomain and promotes chromatin assembly of the host cell factor 1 (HCF-1)-MLL histone methyltransferase complex driving the oncogene expression [82]. This is possible that ATAD2 may promote BCL6 expression via similar mechanisms. The constitutive expression of *ATAD2* and *ATAD5* may have adverse effects, potentially leading to GCDBL formation. This aligns with the expression patterns observed in human tonsil GC B cells and murine GC B cells, where *Atad2* and *Atad5* expression is reduced in plasma B cells compared to GC B cells (Supplementary Figure 5D-E), indicating that timely regulation of *ATAD2* and *ATAD5* may be critical for GC B cell differentiation, and their concurrent downregulation with *BCL6* may be essential for B cell differentiation.

### Defective Migration of GCDBLs in Genotoxic Stress

Finally, we noted a marked upregulation of *RHOA* in HU-treated Raji cells, a p53-dependent gene, which inhibits the RAC1-dependent migration (Figure 7E) [73]. This suggests that GCDBLs exposed to genotoxic stress may exhibit defective migration within the GC compartment due to p53 regulation. Indeed, mouse B cells treated with Nutlin-3a abolished the CXCL13 migration (Figure 7F, G). We propose this reduced migration may limit GCDBLs’ accessibility to LZ compartments, impeding survival feedback from the GC environment and preventing their terminal differentiation. Therefore, p53 regulation may provide quality assurance for GC B cells prior to terminal differentiation by regulating their migration. However, it remains unclear whether wild type p53 maintains the same regulatory influence on *RHOA* expression in GC B cells, which demands future investigation. The p53 driven *RHOA* expression, leading to reduced cell migration and invasion in other cancers is consistent with our finding, placing p53 as a major regulator of cell migration across the cell types [74]. Future work examining the role of wild type p53 will help reveal if it has a similar role in suppressing migration and invasion of GCDBLs, preventing the B-lymphomagenesis during the GC reaction [74].

## Author contribution

SKG designed the original hypothesis, performed experiments, performed data analysis, wrote the original manuscript. JHB critically read and edited the manuscript, provided constructive feedback.

## Acknowledgements

The study is supported by grant number 21K16142 from Japan Science for promotion of sciences (JSPS) to SKG. The Authors thank Dr. Toyohiro Hirai, Dr. Atsuyasu Sato for supporting this work at Kyoto University Hospital. The authors thank Gohar Rehman and Yutaka Hirayama and Dr. Susumu Kohno for assistance with bioinformatics analysis.

## Material and methods

### Cell culture, treatment, treatment with chemicals, cell lysis, immunoblotting

Raji cells were brought from RIKEN brain science institute and Department of Hematology, Kyoto University Hospital. Raji were cultured in RPMI containing 10% FBS, 1% penicillin streptomycin (Wako#161-23181). Hydroxyurea (HU) (Wako Japan# 085-06653) was prepared as per instructions from the manufacturer. For HU treatment of cells, Raji cells were first treated with serum free RPMI culture medium for 18 hours and then replaced with normal RPMI medium for 5 hrs. Then a final concentration of 4 mM HU was maintained in culture medium for 12 hrs. For Thymidine and Nocodazole treatments, Raji cells were treated with 200 nm Thymidine (Nakalai#11100-21, Japan) for 18 hrs. The cells were washed with Normal RPMI culture medium and then replaced with fresh RPMI medium for 5 hrs. After 5 hours, a final concentration of 20 ng/ml of Nocodazole (57591-94, Nakalai, Japan) was added to the culture medium for up to 10 hrs. For Nutlin-3a treatment (S8059, Selleck, Japan) a final concentration of 10 μM was added to directly the cell culture for 12 hrs. For the immunoblotting, Raji cells were lysed as described [27]. Immunoblotting analysis for histone H3K36me3 and β-actin was performed using antibodies (Supplementary Table 2), as per the manufacturer’s instruction.

### Detection of *IGH-MYC* translocation in Raji cells

1000 ng of genomic DNA was used for PCR amplification. Primer used for the long-distance PCR were used as described [6] and are listed in Supplementary Table 1. Primer-start GXL (#R050A, Takara, Japan) was used for the amplification of PCR band with an extension time of 10 minutes per cycle for total 33 cycles.

### Mouse B cell isolation and migration Assay

Normal adult C57BL/6 mice were euthanized, and their spleens were harvested and disrupted in RPMI culture medium. The cell suspension was filtered through a 70 μm filter to obtain a single-cell suspension. Red blood cell lysis was performed according to the manufacturer’s instructions (Biolegend# 420301). Mouse B cells were isolated using the EasySEP Mouse CD19 Positive Selection Kit (Stem Cell Technologies #18954). Total cells were collected by centrifugation at 800 g for 5 minutes at 4 °C. The positively selected cells were cultured in normal RPMI medium for 18 hours before being used in migration assays and treated with Nutlin-3a (Selleck #S8059). Nutlin-3a (10 μM) treatment was applied according to the specified time points. For cell migration assays, transwell inserts were placed in a 12-well plate (Costar, REF #3421) containing RPMI medium (5% FBS) or RPMI medium (5% FBS) supplemented with rmCXCL13 (1000 ng/ml) (BioLegend #583902) for 5 minutes. Subsequently, 1.5 x 10^6^ B cells were added to the inserts and incubated for 4 hours at 37 °C in a CO_2_ chamber. The migrated cells were collected, lysed in 200 μl of CellTiter-Glo® 2.0 (Promega #G9242), and luminescence was measured using a TECAN reader (code # MIC9414). The migration index was calculated by normalizing the luminescence values observed in cells migrated to CXCL13-transwells to the luminescence of cells migrated to control medium transwells.

### Short hairpin RNA knockdown via lentiviral particles

293T cells were cultured until they reached optimal confluency. Transfection was then performed using a mixture of 3 µg of either *sh-Scramble* or *shATAD2* plasmids (*pLKO1-puromycin*), 3 µg of *p-PAX2*, and 3 µg of *pVSVG*, combined in 1 ml of Opti-Mem (Cat# 31985062, Gibco) along with 27 µl of Polyethyleneimine (PEI) solution (1 µg/ml). This mixture was incubated at room temperature for 20 minutes and subsequently added uniformly to the 293T cells. After 24 hours, the culture medium was replaced with fresh medium, and the cells were incubated for an additional 48 hours to allow for lentiviral particle enrichment in the supernatant. The culture medium was then filtered through a 0.2 µM filter, and the resulting lentiviral medium (1 ml) was used to transduce 1 million Raji cells. Following transduction, puromycin (1 µg/ml) or blasticidine (10 µg/ml) was applied for selection. Fresh puromycin or blasticidine was replenished every three days for up to one week to establish stable cell lines.

### FACS analysis

Human CXCR5, CXCR4 and CXCR3 antibodies with respective isotypes was obtained from Biolegend as listed in Supplementary Table 2. For surface staining of human B lymphoma cells, 1 *10^6^ cells were collected and washed twice with FACS buffer (PBS# Nakalai Tasque, 0.1% BSA and 0.1% NaN_3_) and then suspended into a final volume of 100 μl FACS buffer containing the respective antibody or isotype controls. The final dilution of CXCR5, CXCR4 or CXCR3 or respective isotype controls was performed asper the manufacturers instruction. The cells were stained for 15 minutes on ice. The cells were washed twice with FACS buffer and resuspended in 400 μl FACS buffer. A live cell indicator 7-AAD was added just before the flow analysis. Flow cytometry was performed on a FACS Fortessa Flow cytometer (Becton Dickinson, San Jose, USA) and flow cytometry data were analyzed using Flowjo software by total 10,000 events acquired. Data were displayed as two colour plots (SSC-A/APC/PE/BV-421).

### RNA-sequencing computational analyses of RNA-seq, qRT-PCR and identification of p53-DEGs

FASTQ files were generated using Bcl2fastq. Low-quality reads were trimmed out from the FASTQ files using fastp. The reads were mapped on the GRch38 genome reference using HISAT2, and StringTie was used for transcript assembly and quantification. The differential gene expression analysis was performed using R package TCC. Enriched pathways were determined using the GSEA tool available from the Broad Institute website. Gene sets derived from the GO Biological Process ontology were downloaded from the MSigDB database. FX1 treated datasets of Raji cells were taken from published studies [50] (GSE254904). To identify differentially expressed genes associated with p53 signaling, DESeq2 R package was utilized with adjusted p <0.0001 using 2 closely related cohorts of either control, HU and Thy-Noc treated samples [83, 84]. Log2FC values of significantly altered genes was displayed using GraphPad prism. qRT-PCR was performed and analyzed as described before [27].

### Chromatin Immunoprecipitation (ChIP), ChIP-seq library preparation and computational analyses of ChIP-seq

ChIP was performed as previously described [28]. The ChIP-seq library was prepared according to the manufacturer’s instructions using the Thruplex DNA-seq Kit (R400675, Takara Bio, Japan). The prepared ChIP-seq libraries were sequenced by Macrogen, Japan, using the Illumina NovaSeq platform to achieve high-throughput and resolution. FASTQ files were generated using Bcl2fastq. Low-quality reads were trimmed out from the FASTQ files using fastp, and the reads were mapped on the GRch38 genome reference using Bowtie2. MACS2 was used to generate BED files of peak calling for H3K4me3. Visualization of Histone H3K4me3 peaks in control and HU-treated Raji cells was conducted using IGV, along the gene bodies analyzed.

### GEOR datasets and analysis

The RNA-seq and ChIP datasets for this study are available under GEO accession numbers GSE242375 and GSE242936, respectively. The analysis of Raji and DLBCL cell lines treated with FX1 and a BTK inhibitor was conducted using datasets from GEO accession numbers GSE254904 [50] and GSE171763 [85]. Peaks of BCL6 binding in the B-ALL, HepG2, OCI-LY1 and SUDHL4 cell lines were obtained from ChIP atlas (https://chip-atlas.dbcls.jp/data/hg38/target/SRX18259603.1.html). Mouse splenic plasma B cells and GC B cells samples were accessed with GEO accession number GSE60927 [41].

---

**Supplementary Table 1:**
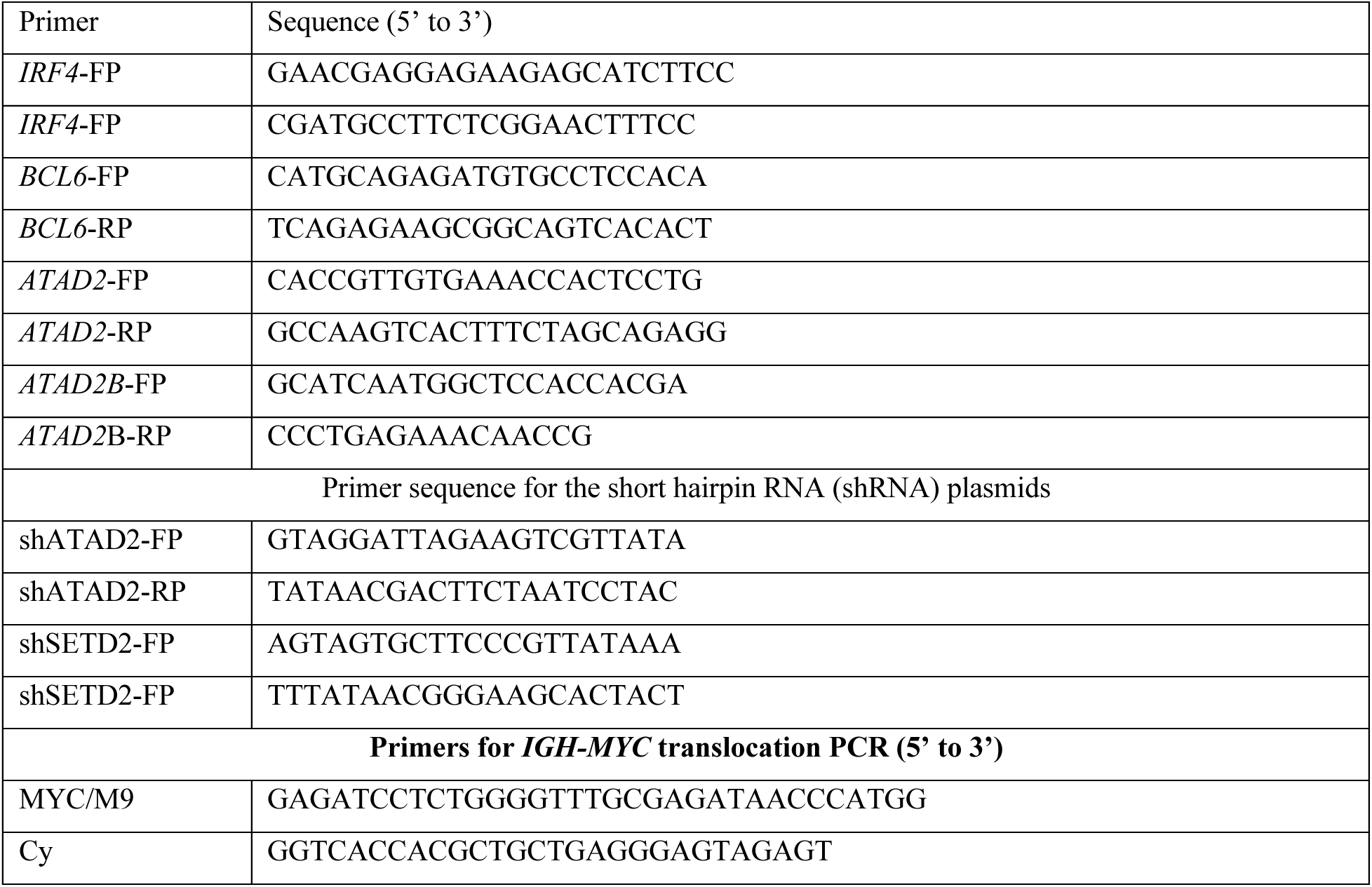

**Supplementary Table 2:**
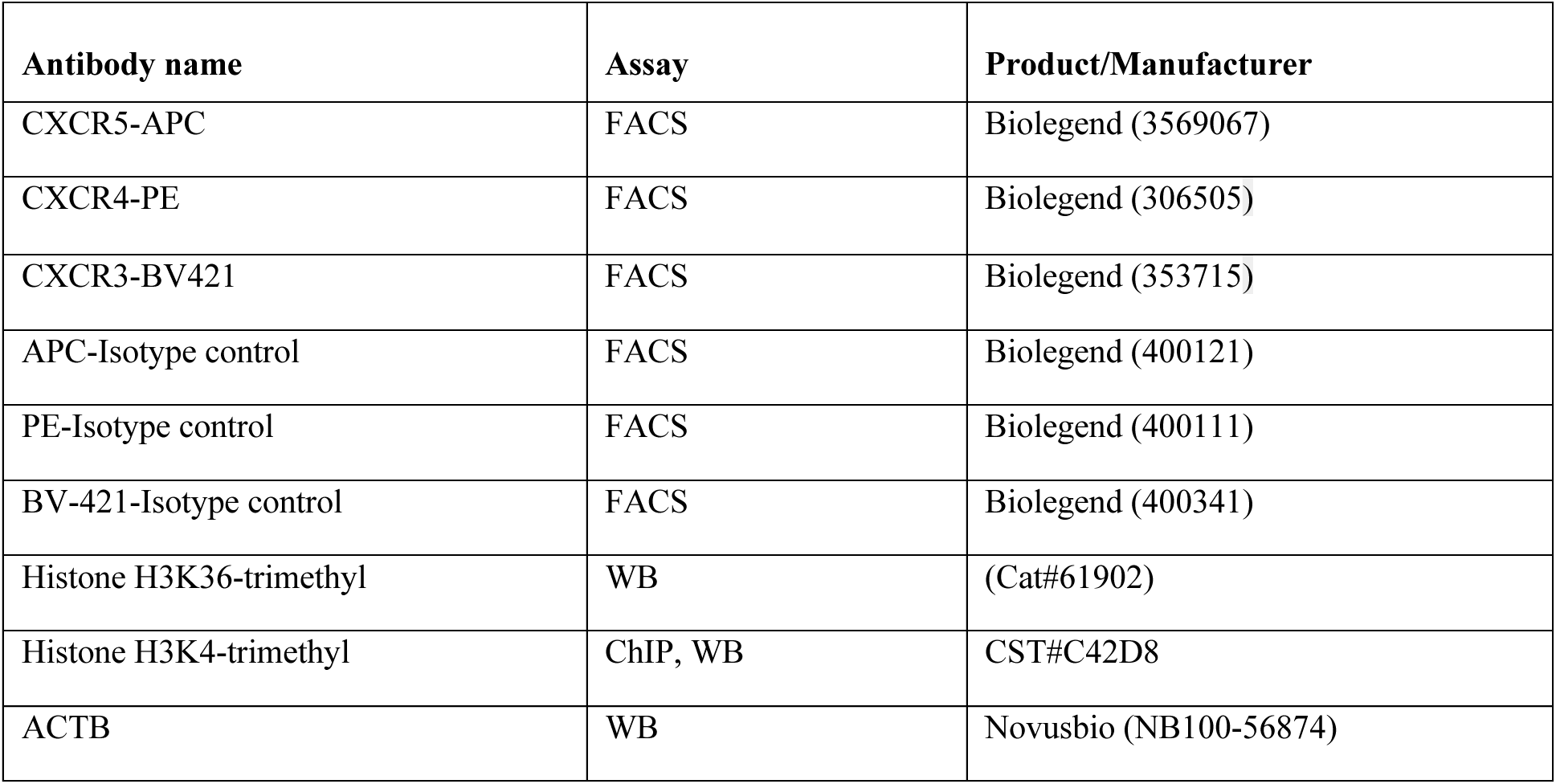

**Supplementary Figure 1.**
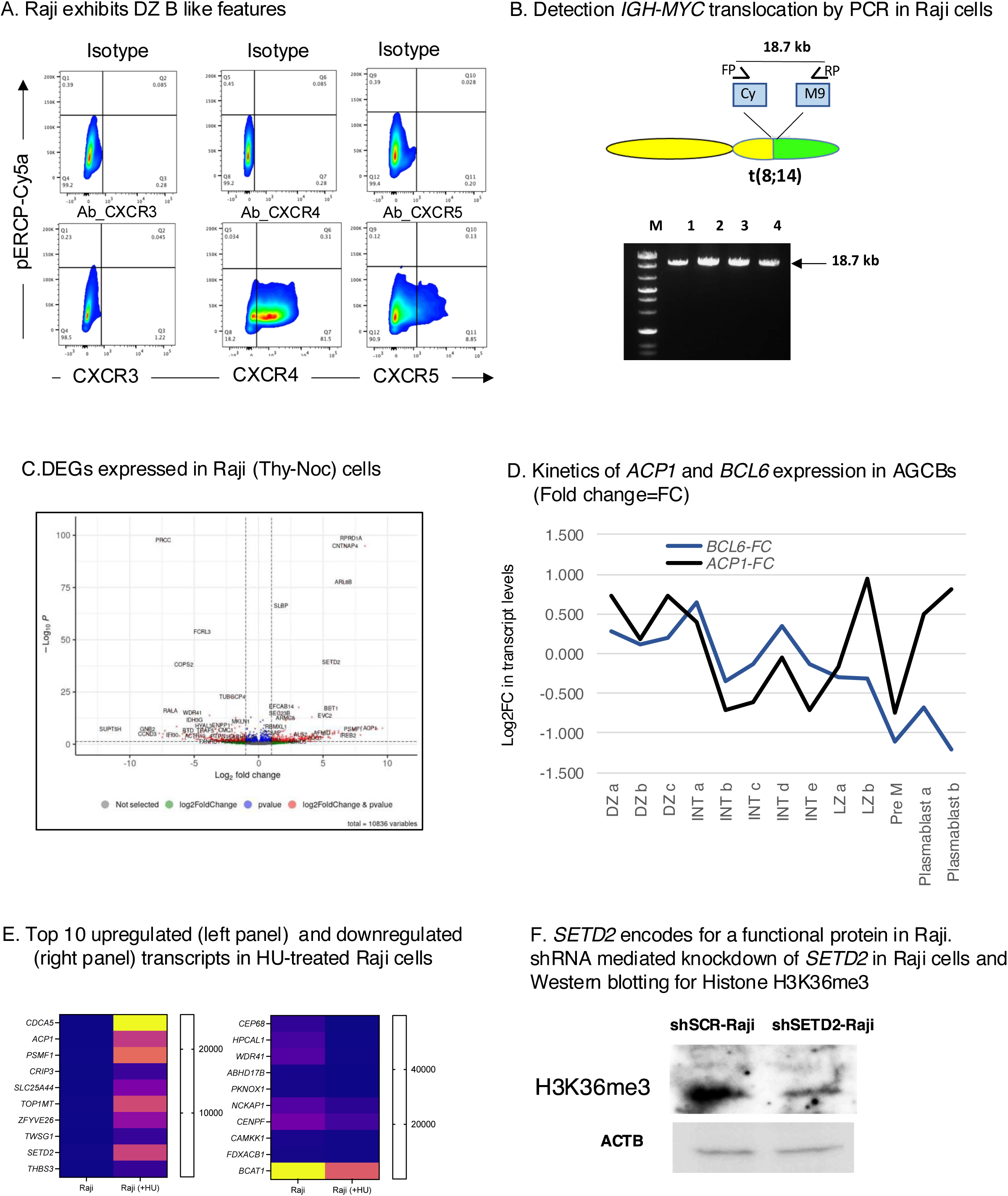
Raji cells exhibit features similar to DZ derived B-lymphomas. **(A)** Raji cells exhibit CXCR5^low^ and CXCR4^hi^ status as surface markers, similar to GC B cells in dark zones. Flow cytometry of surface stained Raji cells with CXCR3, CXCR4, CXCR5 and respective isotype control antibodies **(B)** Detection of *IGH-MYC* translocation in four independent genomic DNA samples from Raji cells. Forward primer (FP) and reverse primer (RP) bind to the Cγ and M9 exons of *MYC*, respectively. PCR was performed with 1000 ng of total DNA for 35 cycles, each cycle lasting 10 minutes, using Primestar GXL (Takara-Bio) **(C)** Volcano plot showing differentially expressed genes between control and Thy-Noc-treated Raji cells. Genes with cutoff p values (p<0.05) are highlighted. The x-axis represents log2 fold changes, and the y-axis represents -log10 adjusted p values. Data represent the cumulative results from three independent groups **(D)** *ACP1* is one among the topmost gene induced in the HU-treated Raji cells. Correlation analysis of *ACP1* and *BCL6* in the DZ sub-populations (DZa, DZb, DZc), Intermediate zone subpopulation (INTa, INTb, INTc, INTd, INTe), LZ sub-populations (LZa, LZb), pre memory (PreM) and plasmablast (plasmablast a, plasmablast b) subpopulations. The Log2 fold change value of each transcript is shown **(E)** list of top 10 up (left panel) and downregulated (right panel) genes in the HU-treated Raji cells. Blue; downregulated, red; upregulated. Data from mean value from three replicated samples are shown. The top 50 significantly altered genes were first sorted by p-values (from smallest to largest) and subsequently ranked based on the log2 fold change in gene expression. **(F)** Western blotting analysis shows reduced Histone H3K36me3 levels in shSETD2-Raji cells than shSCR-Raji cells. Raji cells harbor the functional SETD2 protein. β-actin immunoblotting serving as a loading control.

**Supplementary Figure 2.**
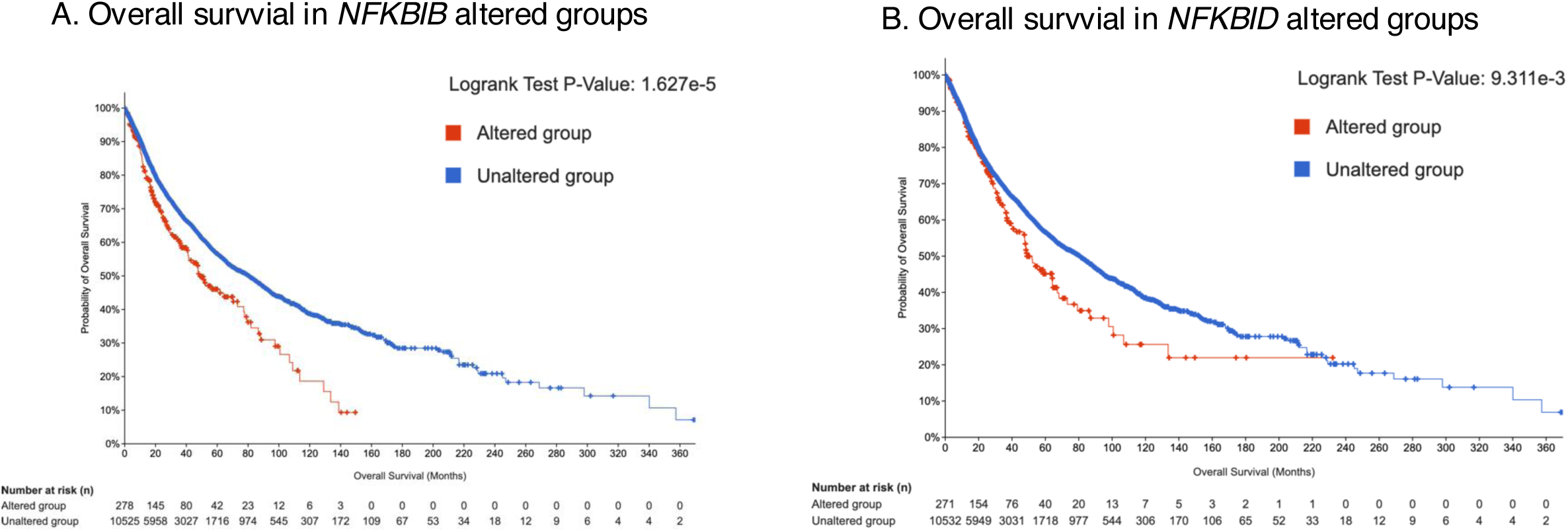
Reduced overall survival of *NFKBIB* and *NFKBID* altered tumors. **(A)** The TCGA datasets showing the overall survival of *NFKBIB* altered groups compared to unaltered groups. Comparison of survival kinetics is initiated with 278 altered groups and 10525 unaltered groups on day 0. p= 1.627e-5 **(B)** The TCGA datasets showing the overall survival of *NFKBID* altered groups compared to unaltered groups. Comparison of survival kinetics of 271 altered groups with 10532 unaltered groups. p= 1.9311e-3.

**Supplementary Figure 3.**
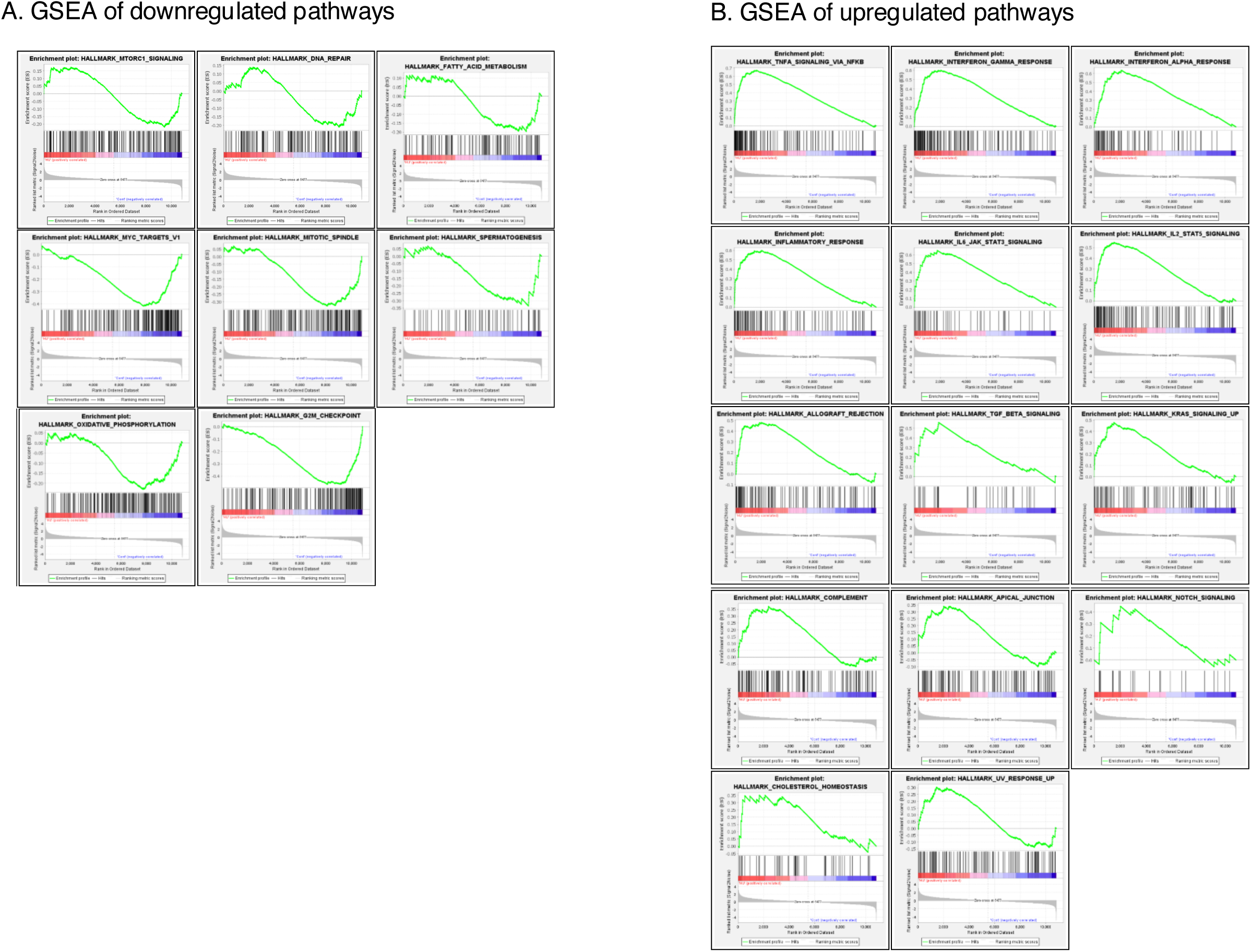
GSEA analysis of upregulated and downregulated pathways in HU-treated Raji cells. **(A)** GSEA analysis of downregulated pathways in HU-treated Raji cells. **(B)** GSEA analysis of upregulated pathways in HU-treated Raji cells. Data are cumulative of three independent replicates. Y axis indicated the enrichment score.

**Supplementary Figure 4.**
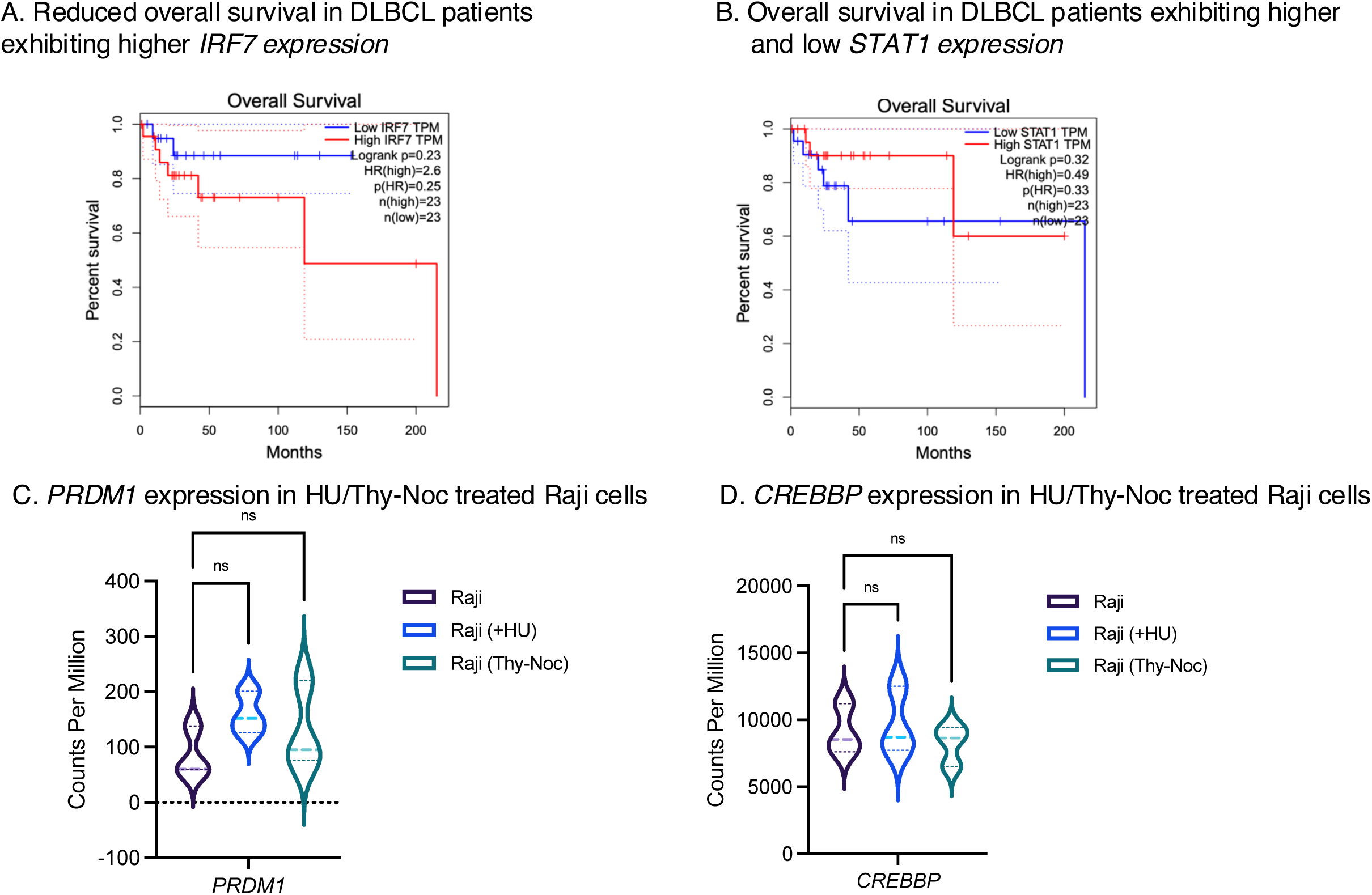
Survival of *IRF7* expressing DLBCLs and expression of *CREBBP* and *PRDM1* in HU and Thy-Noc treated Raji cells. **(A)** DLBCL patients with elevated *IRF7* expression show reduced overall survival compared to those with lower *IRF7* levels, though the decreas is not statistically significant. **(B)** No significant difference in survival is observed between DLBCL patients with high and low *STAT1* expression. DLBCL patient data are obtained from GEPIA database. **(C)** *PRDM1* expression is not altered in Raji (HU) and Raji (Thy-Noc) compared to Raji cells. p=0.2626 (ns) for Raji vs Raji (HU) and p= 0.5582 (ns) for Raji vs Raji (Thy-Noc). One-way Anova; Dunnett’s multiple comparison test. Data are presented as mean ± SEM, n=3. **(D)** *CREBBP* expression is not altered in Raji (HU) and Raji (Thy-Noc) compared to Raji cells. p=0.9302 (ns) for Raji vs Raji (HU) and p= 0.8039 (ns) for Raji vs Raji (Thy-Noc). One-way Anova; Dunnett’s multiple comparison test. Data are presented as mean ± SEM, n=3.

**Supplementary Figure 5.**
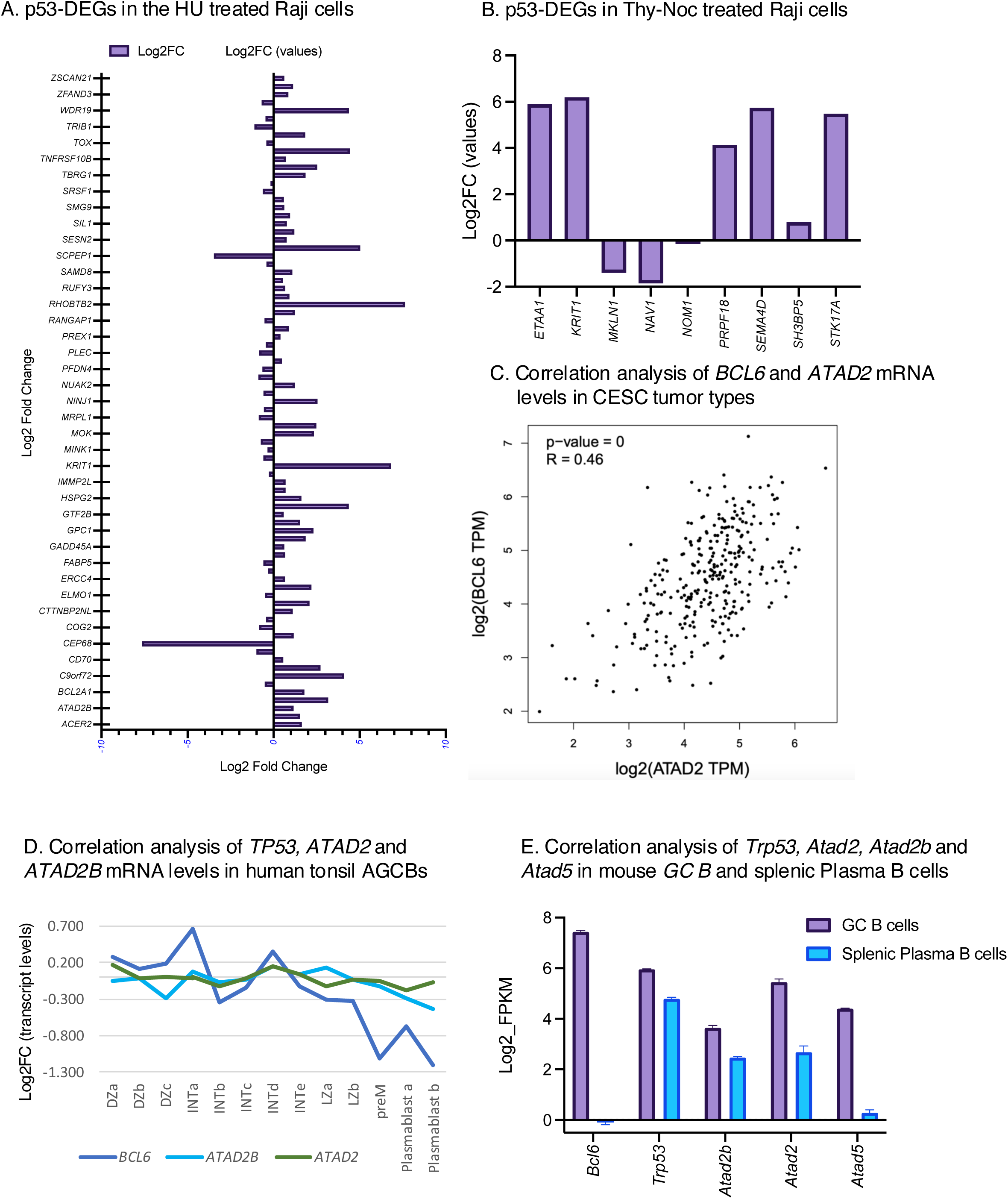
List of p53-DEGs in HU and Thy-Noc Treated Raji Cells. **(A)** Log2 fold change of 81 p53-regulated DEGs in HU-treated Raji cells, with *ATAD2B* positioned as the second-to-last gene. Genes with p < 0.0001 are shown. **(B)** Log2 fold change for nine p53-DEGs in Thy-Noc treated Raji cells. Genes with p < 0.0001 are shown. Data are derived from two closely associate cohorts of HU-treated Raji cells **(C)** Correlation between *ATAD2* and *BCL6* expression across Cervical Squamous Cell Carcinoma (CESC) samples, using GEPIA. R = 0.46 (p =0) **(D)** Correlation of *BCL6*, *ATAD2*, *ATAD2B* transcriptional changes in DZ sub-populations (DZa, DZb, DZc), Intermediate zone subpopulation (INTa, INTb, INTc, INTd, INTe), LZ sub-populations (LZa, LZb), pre memory (PreM) and plasmablast (plasmablast a, plasmablast b) subpopulations of human tonsil AGCBs. Log2 fold change values of each transcript are shown [34]. *BCL6* expression sharply declines from INTd/LZa/LZb to plasmablast b subpopulation. *ATAD2B* expression is declined from LZa to plasmablast b cells **(E)** Expression of *Atad2*, *Atad2b*, and *Atad5* in mouse splenic plasma B cells compared to GC B cells [41] (n=2 for GC B cells, n=3 for splenic plasma B cell samples; GSE60927). *Bcl6* expression is sharply reduced in the plasma B cells*. Atad2, Atad2b* and *Atad5* expression is also reduced in the plasma B cells compared to the GC B cells.

